# Visual mate preference evolution during butterfly speciation is linked to neural processing genes

**DOI:** 10.1101/2020.03.22.002121

**Authors:** Matteo Rossi, Alexander E. Hausmann, Timothy J. Thurman, Stephen H. Montgomery, Riccardo Papa, Chris D. Jiggins, W. Owen McMillan, Richard M. Merrill

## Abstract

Many animal species remain separate not because they fail to produce viable hybrids, but because their individuals “choose” not to mate. However, we still know very little of the genetic mechanisms underlying changes in these mate preference behaviours. *Heliconius* butterflies display bright warning patterns, which they also use to recognize conspecifics. Here, we couple QTL for divergence in visual preference behaviours with population genomic and gene expression analyses of neural tissue (central brain, optic lobes and ommatidia) across development in two sympatric *Heliconius* species. Within a region containing 200 genes, we identify five genes that are strongly associated with divergent visual preferences. Three of these have previously been implicated in key components of neural signalling (specifically an *ionotropic glutamate receptor* and two *regucalcins*), and overall our candidates suggest shifts in behaviour involve changes in visual integration or processing. This would allow preference evolution without altering perception of the wider environment.

---

The evolution and maintenance of new animal species often relies on the emergence of divergent mating preferences^1,2^. Changes in sensory perception or other neural systems must underlie differences in innate behaviours between species, and will ultimately have a genetic basis. Although the significance of behavioural barriers for speciation has been recognized since the Modern Synthesis^3^, we know little of the genes underlying changes in mating preferences, or variation in behaviours across natural populations more broadly^4,5^. Identifying these genes will provide an important route towards understanding how behavioural differences are generated, both during development and across evolutionary time.

Previous studies of isolating preference behaviours have largely been limited to the identification of causal genomic regions, which almost invariably contain many genes^6,7,8^. Only a handful of studies have identified likely candidate genes that contribute to species behavioural preferences. These are largely limited to chemosensory-guided mating preferences^9,10,11^ but see^12^, and have identified changes at chemoreceptor genes. To our knowledge, only two studies – in incipient fish species – have identified candidates for visual preference evolution, albeit indirectly, both suggesting a role for sensory perception mediated by changes in the peripheral visual system (e.g. opsins)^13,14^. Whether or not visual preference evolution generally involves shifts at the sensory periphery, or in downstream processing, remains unknown.

The closely related species *Heliconius melpomene* and *H. cydno* differ in warning patterns, which are both under disruptive selection due to mimicry^15^ and are important mating cues^16^. As a result, these divergent patterns couple ecological and behavioural components of reproductive isolation, which (as predicted by so-called “magic trait” models^17^) is expected to facilitate speciation in the face of gene flow. In central Panama, *H. melpomene* shares the black, red and yellow pattern of its local *Heliconius erato* co-mimic. In contrast, *H. cydno* mimics the black and white patterns of *H. sapho. H. melpomene* and *H. cydno* remain separate largely due to strong assortative mating^18^. Visual preferences for divergent patterns are particularly apparent in males, which strongly prefer to court conspecific females^16,19,20^. Differences in warning pattern between *H. melpomene* and *H. cydno* are largely due to expression differences in just three genes, specifically *optix*^21^, *WntA*^22^ and *cortex*^23^.

Quantitative trait locus (QTL) mapping of *H. melpomene* and *H. cydno* has revealed three genomic regions of major effect that influence the relative time males spend courting red *H. melpomene* or white *H. cydno* females^20^. Notably, the best supported QTL was in the same genomic region as *optix*, the gene responsible for presence and absence of red colour pattern elements in *Heliconius*^21^. Genetic linkage will facilitate speciation by impeding the breakdown of genetic associations between ecological and mating traits^23^. Nevertheless, this QTL, and its associated candidate region, contain hundreds of genes, and the exact genes responsible for differences in preference behaviour are not known.

Here, we first confirm that behavioural QTLs identified previously are associated with variation in male courtship initiation. We then identify genes within the major QTL, which were differentially expressed in the neural tissue (central brain, optic lobes and ommatidia) of *H. melpomene* and *H. cydno*, or have protein coding changes predicted to alter protein function. Out of 200 genes within the QTL region, we identify just five candidates likely to underlie assortative mating behaviours.

## Results

### Chromosome 18 is associated with differences in courtship initiation

Our previous results reveal that QTLs on chromosomes 1, 17 and 18 influence the relative time hybrid males spend courting red *H. melpomene* or white *H. cydno* females^19^. However, the time males spend courting a particular female might depend not only on male attraction, but on the female’s response (and in turn his response to her behaviour). To confirm that these previously reported QTLs influence male approach behaviours (as opposed to other traits that may influence courtship, for example male morphology^25^), we reanalysed our previous data, by explicitly considering whether males initiated courtship towards *H. melpomene, H. cydno* or both types of female during choice trials.

Consistent with our previous analyses^20^, we found that males of both species, and in particular *H. melpomene* males, show a strong preference for females of their own phenotype. F1 and backcross-to-*melpomene* prefer to court *melpomene* females, whereas courtship initiation behaviours segregate in the backcrosses to *cydno* (Figure 1). The QTL on chromosome 1 was retained in our model of initiation behaviours (Supplementary figure 1; *n* = 139, ΔELPD = -13.6 (SE±5.7), *i*.*e*. a change of 2.34 SE units). In contrast, the QTL on chromosome 17 was not retained (*n* = 139, ΔELPD = -2.1 (S.E.±3.0)). Notably, backcrosses-to-*cydno* males heterozygous (*i*.*e*. with a *H. melpomene* allele derived from the F1 father) at the best supported QTL from our previous analysis^20^ (on chromosome 18) initiated courtship towards *H. melpomene* females more frequently than males homozygous for the *H. cydno* allele (Figure 1, bottom left; *n* = 139, ΔELPD: -10.9 (S.E.±5.1), *i*.*e*. a change of 2.14 SE units). Together with previous evidence that male hybrids bearing *H. melpomene* alleles at *optix* prefer to court the artificial models of *H. melpomene* females over those of *H. cydno*^26^, these results suggest that the QTL on chromosome 18 harbours genes for visual attraction behaviours towards females with the red pattern (and that the *H. melpomene* alleles are dominant). As a result, we focused our subsequent analyses on this QTL on chromosome 18 (and also because tight linkage of *optix* allowed us to track the alleles at the preference locus in hybrid crosses). A recent study^27^ also reports no candidate chemosensory genes for reproductive isolation at this QTL region on chromosome 18 (or at any other QTL).

**Figure 1.**
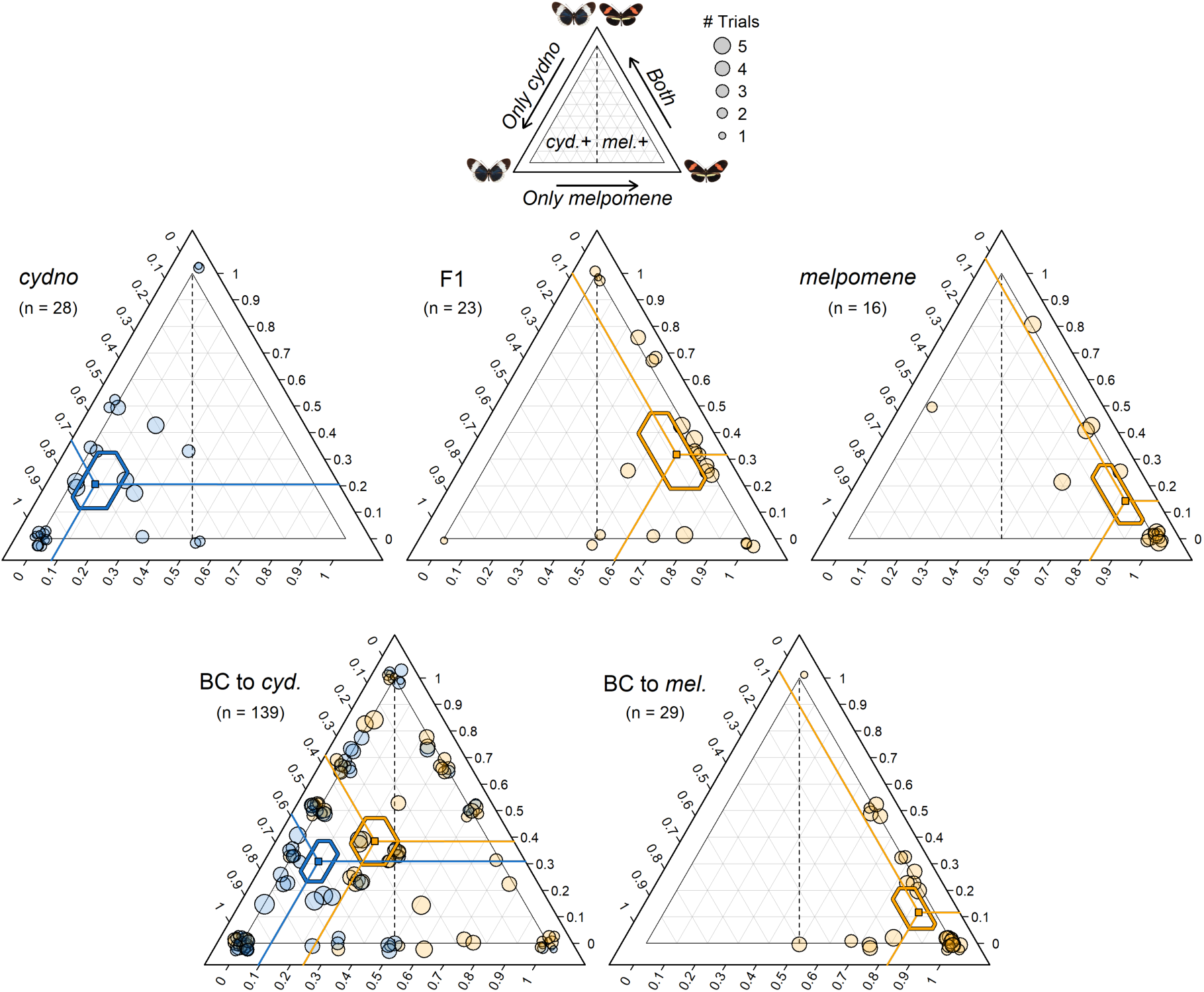
Genotype at the preference QTL on chromosome 18 influences courtship initiation. Ternary plots showing the proportion of 15-minute choice trials in which single male individuals initiated courtship towards *H. melpomene, H. cydno* or both females. Left ternary axis shows the proportion of trials where courtship was initiated towards *H. cydno* female only, bottom axis towards *H. melpomene* female only, and right axis towards both female species. Trials without male response were removed from the dataset. Each point represents a single individual and the location of the point in the triangle is a way of representing these three proportions at once. Lines project the three predicted proportions to corresponding values on the three axes and 95% credibility intervals (CrIs) for these proportions are shown as hexagons. Orange points represent individuals that have inherited at least one *H. melpomene* derived allele at the preference QTL on chromosome 18 (i.e. either *melp*/*melp* or *cyd*/*melp)*; and blue points represent individuals that are homozygous for *H. cydno* alleles at the preference QTL on chromosome 18 (i.e. *cyd*/*cyd*). Point size is scaled to the number of trials in which the male showed a response and a ‘jitter’ function has been applied (leading to some dots being jittered to outside the triangle).

### 27 *genes* within the major QTL are differentially expressed in the brains and eyes of *H. cydno* and *H. melpomene*

We hypothesized that changes in gene regulation that determine differences in visual mate preference behaviours might occur during pupal development (for instance, during visual circuit assembly) or at the adult stage, and must involve changes in the peripheral and/or central nervous system^28^. Therefore, we generated RNA-seq libraries for combined eye and brain tissue, across two pupal stages (around the time of ommochrome pigment deposit and half-way through pupal development) and one adult stage, for *H. melpomene* and *H. cydno* and compared their gene expression levels. Across the QTL region on chromosome 18 (which spans 2.75 Mb and contains 200 genes), we identified 27 genes that show differential expression between *H. melpomene* and *H. cydno*, in at least one of the three developmental stages. These were mostly located within the QTL peak (*i*.*e*. the genomic region with strongest statistical association with male preference) or in close proximity to *optix* (Figure 2). The same genes were frequently differentially expressed across development (Supplementary table 1), with 11 genes being differentially expressed in more than one stage.

**Figure 2.**
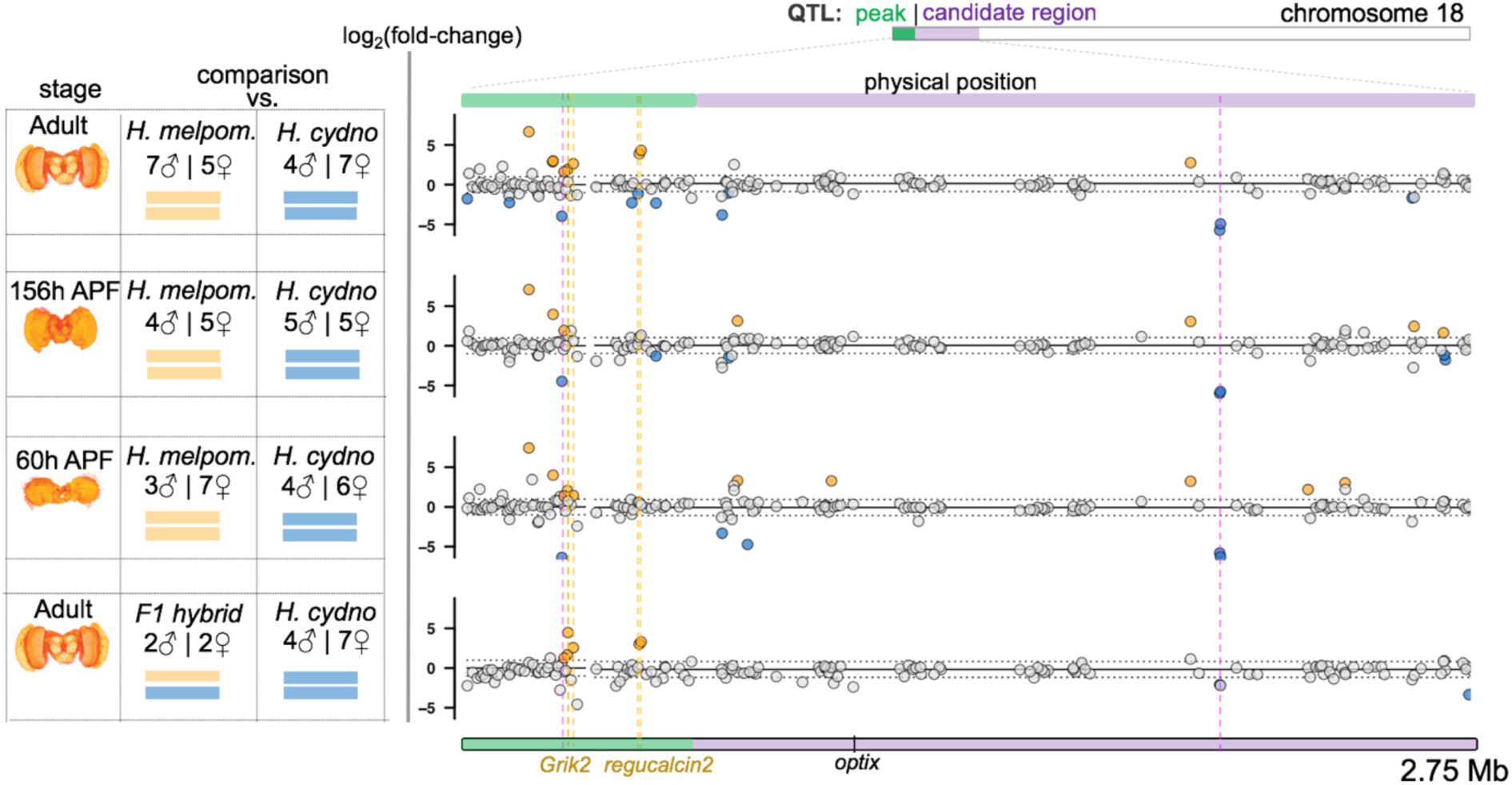
Differential expression at the preference QTL region on chromosome 18. Left: Summary of the comparative transcriptomic analyses with stage, number of samples and chromosome 18 composition. Right: the corresponding results, zooming in on the QTL region on chromosome 18. The *x*-axis represents physical position. The QTL peak, and the rest of the QTL 1.5 LOD candidate region (from^28^) are shown in green and purple, respectively. Points correspond to individual genes, with the *y*-axis indicating the *log*_*2*_(fold-change) for each comparison. The two horizontal dashed lines (at *y*-values of 1 and -1) indicate a 2-fold change in expression. Genes showing a significant 2-fold+ change in expression level between groups are highlighted in orange and blue, where orange indicates higher levels in *H. melpomene* or in the hybrids *cyd*/*melp* (blue if in *H. cydno* – hybrids *cyd*/*cyd*). Vertical dashed lines highlight those genes that are differentially expressed between *H. melpomene* and *H. cydno* AND between *cyd/melp* vs *cyd/cyd* individuals, at the same stage (*Grik2* and *regucalcin2* at the adult stage, *Grik2* at 60h APF). Two genes highlighted by dashed fuchsia vertical lines were excluded because they did not show differential expression, or showed reversal of the fold change when mapping RNA-seq reads to the *H. cydno* genome. Note that *Heliconius* brain (reconstruction) images, added for reference, do not include the eyes (ommatidia and retinal membrane).

The genomic region between the start of chromosome 18 and *optix* (comprising the QTL peak) is highly divergent between *H. melpomene* and *H. cydno*^29^, and divergent gene sequences within this region could also introduce mapping biases of RNA-seq reads. To account for this, we repeated the analysis having mapped to both the *H. melpomene* reference genome^30^ and to a *H. cydno* genome^31^. Generally, we found similar patterns of differential expression when mapping to the *H. cydno* genome (Supplementary figure 2, Supplementary table 2A). Nevertheless, in subsequent analyses we excluded two genes, HMEL034187g1 and HMEL034229g1, which showed reversal of the fold change or did not show differential expression when mapping to the *H. cydno* genome respectively. Results for the QTL on chromosome 1 are reported in the supplementary materials (Supplementary table 1).

### A *regucalcin* and an *ionotropic glutamate receptor* are upregulated in both *H. melpomene* and F1 hybrid males, across independent sequencing experiments

Our previous behavioural experiments suggest that the alleles for the *H. melpomene* behaviour are dominant over the *H. cydno* alleles^20,26^ (Figure 1). Given this pattern of dominance, we predicted that genes underlying variation in male preference to be up- or down-regulated in the brains of both *H. melpomene* and first generation (F1) hybrid males, with respect to *H. cydno*. Of the putative genes differentially expressed between *H. cydno* and *H. melpomene* reported above, only four, within the QTL candidate region, were differentially expressed between the F1 hybrids and *H. cydno* (Figure 2). These included two *regucalcins* (also called *senescence marker proteins-30*: HMEL013552g1, HMEL034199g1), an *ionotropic glutamate receptor* (HMEL009992g4), which is a putative ortholog of *Grik2*, and one gene with no annotated function (HMEL009992g1). We obtained the same results regardless of whether we considered both males and females together, or males alone. Further inspection of spliced mRNA-reads indicated that the two annotated *regucalcins* were in fact a single gene (from now on referred to as *regucalcin2*). This was also the case for the *ionotropic glutamate receptor* and the gene with no annotated function (from now on referred to as *Grik2*).

To ensure that the apparent fragmentation of a few gene models in the *H. melpomene* (Hmel2.5) annotation^32^ did not introduce inaccuracies in estimates of differential gene expression, we produced a new transcript-based annotation of the *melpomene* genome using Cufflinks’s RABT^33^. Repeating all comparative transcriptomic analyses using this new annotation (where exons previously considered to be distinct genes are now assigned correctly to single genes), we confirmed that both *regucalcin2* and *Grik2* were differentially expressed in both species and hybrids comparisons. Finally, to determine whether differential expression of these genes was repeatable across sequencing experiments, we generated RNA-seq libraries for additional *H. melpomene, H. cydno and* F1 hybrids adult brains, sampled five years after our initial tissue collection and sequenced independently. Differential expression of *Grik2* and *regucalcin2* is consistently detected both between *H. melpomene* and *H. cydno* and between F1 hybrids and *H. cydno*, both when these datasets are analysed separately, and when combined.

### Differential expression of *Grik2* and *regucalcin2* in adults is likely due to cis-regulatory effects

Causal changes in gene regulation underlying phenotypic variation associated with the QTL must result from *cis*-rather than *trans-*regulation. In other words, if changes in expression of *Grik2* and *regucalcin2* account for the observed shifts in behaviour associated with the QTL these must be due to changes within the *cis*-regulatory regions of the genes themselves (as opposed to of other *trans*-acting genes elsewhere in the genome); and these causal mutations should be within the QTL region on chromosome 18. To determine whether differences in gene expression levels between parental species were due to *cis*- or *trans*-regulatory changes, we conducted allele specific expression (ASE) analyses in adult F1 hybrids (from both sequencing batches). In F1 hybrids, both parental alleles are exposed to the same *trans*-environment, and consequently *trans*-acting factors will act on alleles derived from each species equally (unless there is a change in the *cis*-regulatory regions of the respective alleles). Therefore, differences in allele specific expression indicate changes in *cis*-regulatory regions^34^. For both candidate genes (*Grik2* and *regucalcin2*) the *H. melpomene* allele was significantly more highly expressed relative to the *H. cydno* allele (p<0.001, Wald test), suggesting *cis*-regulatory effects (Figure 3).

**Figure 3.**
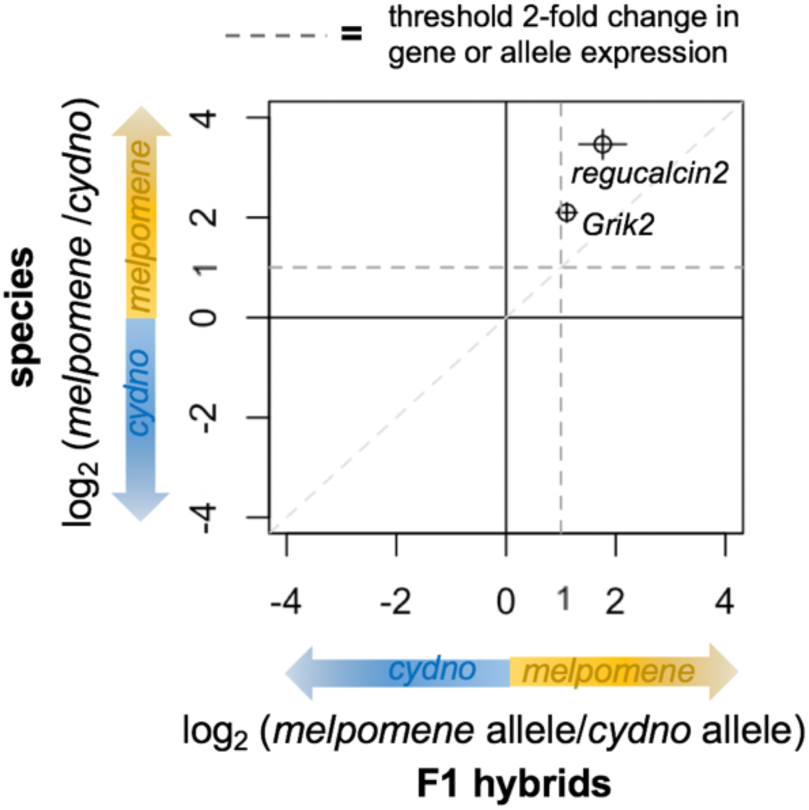
*Grik2* and *regucalcin2* show evidence of allele specific expression. Points indicate the value, and bars the standard error, of the (base 2) logarithmic fold change in expression between parental species (vertical) and the alleles in F1 hybrids (horizontal), for candidate genes (as defined in the transcript-guided annotation). Dashed lines indicate the threshold for a 2-fold change in expression for the genes in the species (horizontal), and for the alleles in the hybrids (vertical). Both genes seem to be regulated by a combination of *cis*- *and trans*-acting factors, rather than *cis*-acting factors alone (which would be indicated by y=x).

### *Grik2* is differentially expressed in hybrids that essentially differ only for allelic composition at the behavioural QTL region

In order to study the specific effects that *H. melpomene* derived alleles at the QTL on chromosome 18 had on gene-expression, we introgressed this region into a *H. cydno* background through multiple backcrosses (crossing design in Supplementary figure 3). We wanted to investigate whether differences at this QTL regulated expression of any specific genetic pathway during development, and more generally what changes in genome-wide transcription were observed in hybrids differing (mostly) just at this QTL region. We compared 6 *cyd/melp* vs. 10 *cyd*/*cyd* (at the QTL region on chromosome 18) BC3 hybrids sampled at 156 hours after pupal formation (APF) (Supplementary figure 5A), and 8 *cyd/melp* vs. 9 *cyd*/*cyd* for those at 60h APF.

Across the entire genome, only 23 genes at 156h APF and 29 genes at 60h APF were differentially expressed between third-generation backcross hybrids with *cyd*/*cyd* and *cyd/melp* genotypes at the QTL. Within the QTL candidate region, *Grik2* was the only gene detected as differentially expressed between species and hybrids at these pupal stages (at 60h APF, Supplementary figure 4). Of the remaining 10 genes that were differentially expressed between both species and backcross hybrids at either stage, seven were located within the introgressed region (0 - 6.3 Mb, Supplementary table S3), and it seems most likely that these are regulated by introgressed *H. melpomene cis-*acting elements. We discuss the possibility of a *cis*-regulatory element within the QTL candidate region acting on a gene outside the QTL in the Supplementary information.

**Figure 4.**
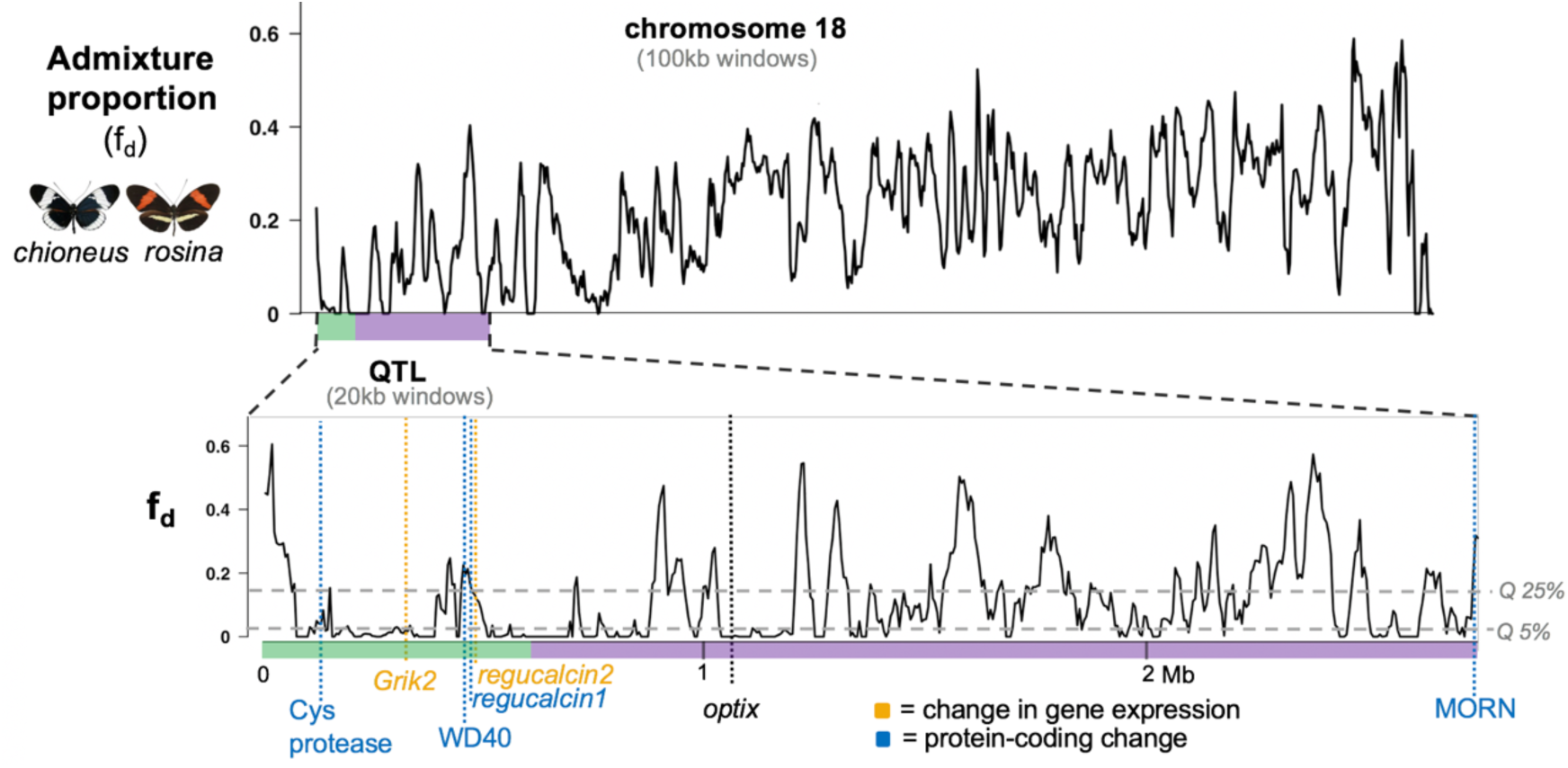
Gene flow at the QTL region for behaviour. Admixture proportion (*f*_d_) values estimated in overlapping 100kb (top) and 20kb (bottom) windows for chromosome 18 (top) and the QTL region (bottom) between *H. melpomene rosin*a and *H. cydno chioneus*, with candidate genes positions highlighted by a vertical dashed line, and *optix* location displayed for reference. The *x*-axis represents physical position, the *y*-axis indicates the *f*_d_ value. *f*_d_ values close to zero indicate that the proportion of shared derived alleles, and consequently gene flow, between *H. melpomene* and *H. cydno* is small (or zero), implying localized selection against foreign alleles that introgress between the two species. 5% and 25% quantile of the *f*_d_ distribution are indicated by horizontal grey dotted lines.

It is possible that causative loci (e.g. expressed RNA or protein factors) within the QTL could act on other genes in *trans* (both on chromosome 18 and elsewhere in the genome), and identifying these could provide insight into the mode of action of causative genetic elements. Three genes differentially expressed both in species and backcross hybrid comparisons, and located outside of introgressed regions, including HMEL014795g1 and HMEL015842g1 (at 10.2 Mb and 13.5 Mb on chromosome 18, respectively) and HMEL030024g1 (on chromosome 1) might be considered good candidates for *trans*-regulation. However, in our backcrosses, the region on chromosome 18 introgressed from *H. melpomene* into a *H*. cydno background extends ∼3.6 Mb beyond the QTL candidate region, making it difficult to determine whether these genes are regulated by loci associated with variation in behaviour. Furthermore, no genetic pathway was enriched for gene expression differences between these hybrids at either pupal stage (PANTHER enrichment test^35^), suggesting that overall this QTL harbours a few, modular changes in gene regulation in the developing brain/eyes of *H. cydno* and *H. melpomene*, or at least not impacting on other genes in *trans* with a clear mode of action.

To verify that differential expression of candidate genes within the QTL region is driven by *H. melpomene* alleles on chromosome 18 and not by other *H. melpomene* alleles at *trans*-acting genes on other chromosomes, we compared gene expression levels between hybrids carrying *cyd/melp* vs. *cyd/cyd* regions on chromosomes chr1, chr4 and chr15, chr20 (Supplementary figure 5A). In these comparisons, there was no signal of differential expression on chromosome 18. This supports the *cis*-regulatory activity of the *melpomene* allele of candidate genes on chromosome 18. To test this further, we conducted another allele specific expression (ASE) study in the BC3 hybrids, which suggested *trans*-only regulatory effects for *Grik2* at these pupal stages, and *cis*-regulatory effects for *regucalcin2* at 60h APF (p<0.038, Wald test) (Supplementary figure 6). Since causal gene/s might exert an effect on behaviour due to their action during development or in adult form, and this action might in turn be differently (*cis*-vs *trans*-) regulated, we still considered both genes as strong candidates.

### 4 genes with protein-coding substitutions within the QTL candidate region have predicted effects on protein function

Because shifts in behavioural phenotypes could be due to changes in protein-coding regions, we additionally considered protein-coding substitutions between *H. melpomene* and *H. cydno*. Overall, we found 152 protein-coding substitutions, spanning 54 of the 200 genes across the entire QTL candidate region. We then studied whether these variants were predicted to have non-neutral effects on protein function with PROVEAN^36^. The PROVEAN algorithm predicts the functional effect of protein sequence variations based on how they affect alignments to different homologous protein sequences. We found 4 genes with such predicted effects (PROVEAN score < -2.5): Specifically, a *WD40*-repeat domain containing protein (HMEL013551g3), a *cysteine protease* (HMEL009684g2), a *MORN* motif containing protein (HMEL006660g1), and another *regucalcin* (HMEL013551g4) adjacent to, but distinct from, that found to be differentially expressed above (from now on referred to as *regucalcin1*).

### Candidate genes occur in regions with reduced gene flow

Of our six candidate genes for preference behaviours that contribute to reproductive isolation between *H. cydno* and *H. melpomene* (*regucalcin2, Grik2* and the four genes with protein coding modifications), five are found within the QTL peak (Figure 4). Genetic changes causing reproductive isolation between populations are expected to reduce localized gene flow in their genomes. Therefore, we compared the position of our candidate genes to estimated levels of admixture proportions (*f*_d_)^37^ between *H. melpomene* and *H. cydno* across the QTL candidate region^38^. We found that candidate genes were located in genomic regions with low *f*_d_ values (Figure 4), suggesting localized resistance to gene flow between *H. melpomene* and *H. cydno* at these genes and their putative *cis-*regulatory regions.

## Discussion

Behavioural isolation is frequently implicated in the formation of new species^1,39^, and often involves the correlated evolution of both mating cues and mating preference. Here we have analysed a genomic region in a pair of closely related sympatric butterflies, *H. cydno* and *H. melpomene*, that contains genes for divergence in both an ecologically relevant mating cue and the corresponding preference. Physical linkage between ecological and mating traits will facilitate speciation by allowing different barriers to act in concert to restrict gene flow^40,41^. Although the genes underlying changes in the warning pattern cue in *Heliconius* are well characterized^21,22,23,42^ (e.g. *optix*), those underlying the corresponding shift in behaviour have not previously been identified^20,43,44^. We have pinpointed a small number of genes that fall within the QTL peak, which show either expression (*regucalcin2* and *Grik2*) or protein coding differences (HMEL013551g3, HMEL009684g2, and *regucalcin1*) and fall within a region of reduced admixture, that are strong candidates for modulating mating behaviour.

Two broad neural mechanisms could underlie the evolution of divergent visual preferences, involving changes in either i) detection at the sensory periphery or ii) the processing and/or integration of visual information. Although *H. melpomene* and *H. cydno* have the same retinal mosaics/class of photoreceptors^45^, spectral sensitivity in the *Heliconius* eyes could be altered by filtering pigments^46^, or other physiological processes taking place at the photoreceptors/sensory periphery, eventually shifting sensitivity towards different wavelengths (and possibly colour patterns). It has previously been hypothesized that the gene regulatory networks for ommochrome deposition in the *Heliconius* eyes might have been co-opted in the wings^47^, where *optix* plays a central role, and therefore that *optix* might play a role in eye pigmentation in *Heliconius*. However, the protein product of *optix* has not been detected in pupal or adult retinas of various *Heliconius* species tested^48^, and therefore has no obvious link to ommochrome deposition in the eyes. More generally, the underlying evolutionary mechanism is unlikely to involve detection at photoreceptors, as this would probably have a broad effect on downstream processing^2^ and alter the visual perception of the animal’s wider environment.

The second mechanism, involving changes in the processing, and/or integration, of visual information, could act through an alteration of neuronal activity or connectivity. For instance, different levels of gene expression in conserved neural circuits between *H. melpomene* and *H. cydno* may affect overall synaptic weighting and determine whether a signal (e.g. colour and motion) elicits a motor pattern (response towards a female) or not. Consistent with this scenario, the composition of ionotropic receptors at post-synapses is a key modulator of synaptic transmission^49^, implicating *Grik2*. Differential expression of ionotropic receptors is also associated with variation in female preference behaviours in fish^50,51.52^, raising the possibility that ion channels might provide a likely route to modulate mate preferences across taxa more broadly. *Regucalcins* are involved in calcium signalling^53^, which regulates synaptic excitability and plasticity^54^, and has an important role in axon guidance^55^ (albeit alongside additional roles across a broad range of biological processes), making the two *regucalcins* we identify strong candidates for behaviour.

Changes in the regulation of genes with pleiotropic effects are likely to be less detrimental compared to changes in their protein-coding sequences^56^ (although emerging evidence has begun to suggest that enhancer/repressor elements may be more pleiotropic than previously thought^57,58^). Furthermore, there is considerable evolutionary potential in the co-option of transcription factors/networks^56^ that regulate neural patterning or neuron-type activity, possibly resulting in novel adaptive expression patterns. In line with this, *regucalcin2* and *Grik2*, which are differentially expressed in the eyes and brain in both our species and hybrid comparisons, are likely to be involved in multi-functional processes, such as calcium signalling and ion transport, and likely have pleiotropic alleles. We also found evidence of *cis*-regulatory effects for both genes, which would be required of the causal genetic change within the QTL, if it were to be in gene regulation.

Neither *regucalcin2* or *Grik2* show male-biased gene expression, which might be expected of candidate genes for a behaviour that is evident only in males. It is possible that the lack of sex-biased expression indicates that visual cues are similarly important in female mating preference in this species pair. Indeed, Chouteau *et al*^59^ report that female preferences contribute to *disassortative* mating between colour pattern morphs of *H. numata*, and it is possible that in *H. cydno* and *H. melpomene* females share the same genetic basis for colour pattern-based discrimination as males. Alternatively, if visual preference behaviour is restricted to males, changes in gene expression may be integrated differently in the female and male nervous systems. The role of female preference in *Heliconius* mate choice remains poorly understood. Although emerging data suggests that female choice does contribute to reproductive isolation between *melpomene-cydno* clade taxa^60,61^, it remains to be tested if there is a strong visual component to this preference similar to that observed in males.

Despite expectations that non-coding, regulatory loci may provide a flexible route to divergent mating preferences, we also found substitutions in coding regions at the QTL, which are predicted to have an effect on protein functioning and therefore remain strong candidates. These genes include *regucalcin1*, which is distinct from, but located next to, *regucalcin2* (which is differentially expressed). Notably, the eye transcript of *regucalcin1* was recently characterized as fast-evolving across *Heliconius* species^62^. Other candidates include a *cysteine protease*, which functions in protein degradation, and might be linked to behaviour for example through degradation of neurotransmitters, a *MORN* motif containing protein (function unknown), and a *WD40* containing protein. *WD-repeat* containing proteins have been implicated in a wide array of functions ranging from signal transduction to apoptosis (https://www.ebi.ac.uk/interpro).

Although preference for red colouration and the *optix* gene are tightly linked, we find no evidence that *optix* is differentially expressed in the eyes or brains of our two species. It is also not located within the QTL peak (and it contains no non-synonymous changes in protein coding regions^21^). It seems unlikely therefore that changes in cue and preference are pleiotropic effects of the same allele. More generally, although we have pinpointed the strongest candidates yet identified for assortative mating behaviours in *Heliconius*, it is possible that actual causal changes in gene regulation are restricted to developmental stages other those sampled, or restricted to a few neuronal populations not detected with transcriptomic data from eyes and whole brain tissue. Nonetheless, by sampling at two pupal stages (around the time of *optix* expression/ommochrome pigment deposit in the wing/eye and halfway through pupal development) and at the adult stage, we should have captured important transitions for the behavioural programming of the two species.

Work in the past decade has shown that complex innate behavioural differences between species can be encoded in relatively few genetic modules^63,64^, but very few studies^65,66,67^ have identified specific genes underlying behavioural evolution. In particular, traditional laboratory organisms continue to provide important insights into the evolution and genetics of behaviour^,28,66,68^, however, comparative approaches are required to determine if developmental principles can be broadly applied, and also to incorporate a wider range of phenotypic variation and sensory modalities. The challenge now is to increase the resolution of studies in non-traditional systems, in order to link individual genetic elements to behaviours, *and* the sensory and/or neurological structures through which they are mediated. In this light, we have identified a small handful of strong candidate genes associated with the evolution of visual mate preference behaviours in *Heliconius*. These genes are in tight physical linkage with the locus for the corresponding shifts in an ecologically relevant mating cue, providing an important opportunity to investigate the build-up of genetic barriers crucial to speciation. The candidate genes identified seem more likely to alter visual processing or integration, rather than detection at photoreceptors, consistent with permitting changes in mate preference without altering perception of the animal’s wider environment.

## Methods

### Courtship initiation analyses

Butterfly rearing, crossing design and genotyping are described in detail elsewhere^20^. In brief, we assayed male preference behaviours for *H. melpomene, H. cydno*, their first generation (F1) hybrids and backcross to hybrids to both parental species in standardized choice trials. Males were introduced into outdoor experimental cages (1×1×2m) with a virgin female of each species and courtship behaviours recorded. Whenever possible, trials were repeated for each male (median = 5 trials). To determine whether previously identified QTLs for courtship time contribute to variation in courtship *initiation* behaviours, we performed a *post-hoc* analysis using categorical models in a Bayesian framework with a multinomial error structure, using the R package *brms*. All models were run under default priors (non- or very weakly informative). In contrast to our previous analysis^20^, in which we considered the number of minutes (*i*.*e*. time) for which courtship was directed towards *H. cydno* or *H. melpomene* females, here the response variable was number of trials in which male courtship was *initiated* towards *H. cydno* females only, *H. melpomene females* only, or both female types (hereafter referred to as “*initiation”*). Across males the median number of trials with a response was 3. Using backcross-to-*cydno* males only, we fitted *initiation* as a response variable to genotype (*cyd*/*cyd* or *cyd*/*melp*) at each QTL, which were included as separate fixed effects. Individual ID was fitted as random factor. To test the effect of each QTL on male *initiation*, we compared the saturated model incorporating all three QTL with reduced models excluding each QTL in turn, using approximate leave-one-out (LOO) cross-validation^69^ as implemented in *brms*, and based on expected log pointwise predictive density (ELPD). Normal distribution of ELPD can be a straightforward approximation given our large samples sizes (n=139)^69^. Therefore, we considered an absolute value of ELPD greater than 1.96 units of its standard error as indicative of the reduced model being less-informative than the saturated model (95% confidence). Males that did not initiate courtship to any female across trials were excluded from analyses, resulting in a dataset of 139 males, from a total of 146 backcross males for which we had genotype data. Finally, we extracted predictors and credibility intervals for backcross males with differing genotypes from the minimum adequate model. Credibility intervals for *H. melpomene, H. cydno*, F1 hybrid and backcross to *melpomene* males displayed in Figure 1 were generated following the same procedures. Raw data and analysis code are available in the following github repository: https://github.com/SpeciationBehaviour/neural_genes_heliconius.git

### Butterfly collection, rearing and crossing design for expression analyses

Wild *H. melpomene rosina* and *H. cydno chioneus* individuals were caught along Pipeline Road near Gamboa, Panama, in the Soberania National Park, and used to establish stocks at the Smithsonian Tropical Research Institute insectaries in Gamboa. Butterflies were reared in common garden conditions, in 2×2×2m cages, and provided with fresh *Psiguria* flowers and 10% sugar solution. Larvae were reared on fresh *Passiflora* shoots/leaves until pupation. *H*. cydno, *H. melpomene* and hybrid individuals used for RNA-seq (see below) were reared concurrently and under the same conditions. F1 hybrids were obtained by crossing a wild-caught *H. m. rosina* male to an insectary-bred virgin *H. c. chioneus* female.

Third-generation backcross hybrids (BC3) were generated by outcrossing a hybrid male with a red forewing band (crossing design shown in Supplementary figure 3) to virgin *H. cydno* females, over three generations. The peak of the behavioural QTL reported previously^20^ on chromosome 18 (at 0cM) is in very tight linkage with the *optix* colour pattern locus (at 1.2cM), which controls for the presence and absence of the red forewing band seen in *H. melpomene rosina*. Presence of the red forewing band is dominant over its absence so that segregation of the red band can be used to infer genotype at the *optix* locus. Specifically, hybrid individuals with a red forewing band are heterozygotes for *H. melpomene/H. cydno* alleles at the *optix* locus, whereas individuals lacking the red band are homozygous for the *H. cydno* allele. Due to the tight linkage we expected little recombination between *optix* and QTL peak even after three generations of introgression, allowing us to infer genotype at the preference-*optix* locus (which we confirmed with genetic data, see below).

### Tissue dissection, RNA extraction and mRNA sequencing

Eye (ommatidia and retinal membrane) and brain tissue (central brain and optic lobes) were dissected out of the head capsule (as a single combined tissue) in cold (4 °C) 0.01M PBS solution, at two pupal stages: 60 hours after pupal formation (60h APF) and 156h APF; and in adults aged 9 - 13 days in 2013. We sampled adults at around 10 days of age because by this stage males are mature and frequently court females^70^. Adult males and females sampled were sexually naive. We decided to sample at 60h APF because this is the developmental stage at which *optix* is expressed in the wing, so we hypothesized that it might had also been when *optix* is expressed in the brain. We sampled at 156h APF as a putative stage halfway through pupal development, and at this stage most of the major neural connections have just been established in the *Heliconius* brain (Stephen Montgomery, unpublished data).

Tissues were stored in RNAlater at 4 °C for 24 hours, and subsequently at -20 °C, until RNA extraction. Total RNA was extracted using TRIzol Reagent (Thermo Fisher, Waltham, MA, USA) and a RNeasy Mini kit (Qiagen, Valencia, CA, USA). Samples were treated with DNase I (Ambion, Darmstadt, Germany). Integrity of total RNA was checked either on an agarose gel or using an Agilent Bioanalyzer 2100 (Agilent Technologies, Santa Clara, CA, USA). RNA concentration was measured on a Nanodrop spectrophotomer. Illumina TruSeq RNA-seq libraries were prepared and sequenced at Edinburgh Genomics (Edinburgh, UK) with 100 bp paired-end reads (in 2014). To avoid lane effects the distribution of the species samples was randomized on the sequencing platform. More detailed information about individuals and sequencing yields can be found in the Supplementary dataset. In 2019, we independently sampled a further 5 *H. melpomene, 5 H. cydno*, and 6 F1 hybrids males, of 10 days of age. These F1 hybrids were generated by crossing a wild caught *H. c. chioneus* male to an insectary-bred virgin *H. m. rosina* female. Briefly, for these, RNA was extracted similarly with Trizol Reagent and a PureLink RNA Mini Kit, with PureLink DNase digestion on column (Thermo Fisher, Waltham, MA, USA). Illumina 150bp paired-end RNA-seq libraries were prepared and sequenced at Novogene (Hong Kong, China).

### RNA-seq read mapping and differential gene expression analyses

After a quality control of RNA-seq reads with FastQC, we trimmed adaptor and low-quality bases using TrimGalore v.0.4.4 (https://www.bioinformatics.babraham.ac.uk/projects/). RNA-seq reads were mapped to the *H. melpomene* 2.5 genome assembly^30^/annotation^32^ using STAR v.2.4.2a^71^ in 2-pass mode. We only kept reads that mapped in ‘proper pairs’ using Samtools^72^. The number of reads mapping to each gene were estimated with HTseq v. 0.9.1^73^ with model “union”, thus excluding ambiguously mapped reads. Differential gene expression analyses between species/hybrids were conducted in DESeq2^74^, including sequencing batch as random factor when comparing species samples from both datasets. We considered only those genes showing a 2-fold change in expression level, and at adjusted (false discovery rate 5%) p-values < 0.05, to be differentially expressed, to exclude expression differences caused by known differences in brain morphology^75^, that in this species pair are clustered in the visual system^76^, although there is no evidence that visual mate preference is linked to these divergent brain morphologies.

### Sexing pupae

In all DESeq2 analyses, sex was included as a random factor. To sex pupae, we first marked duplicate RNA mapped reads with Picard (https://broadinstitute.github.io/picard/), and used GATK 3.8^77^ to split uniquely mapped reads into exon segments and trim sequences overhanging the intronic regions. We then used Haplotype Caller on each individual, using calling and filtering parameters according to the GATK Best Practices for variant calling on RNA-seq data. The sex of pupal samples was inferred from the proportion of heterozygous (biallelic) SNPs using the R package SNPstats. Males (ZZ) were expected to have >> 0% heterozygous sites, whereas females (ZW) to have 0%. Z-linked heterozygosity of the pupal samples (Supplementary table 4) were in line with expectations (either ∼ 0 for females or an order of magnitude higher for males), and matched heterozygosity of either adult males or females, for which the sex was determined from external morphology.

### Inference of gene function and transcript-based annotation

Biological functions of annotated genes were inferred with InterProScan v5^78^, using the corresponding Hmel2.5 predicted protein sequences. InterProScan uses different databases like InterPro, Pfam, PANTHER, and others, to infer functional protein domains and motifs (based on homology). To study whether specific biological functions were enriched among genes showing differential expression among hybrid types, we conducted the PANTHER enrichment test^35^ (with Bonferroni correction for multiple testing) using *Drosophila melanogaster* as the reference gene function database.

Upon detailed inspection of the mapping coverage of spliced RNA-seq reads to the Hmel2.5 gene annotation, we noticed that some gene models were fragmented, namely, a few exons that appeared to be spliced together were incorrectly considered distinct genes. To check that this did not introduced inaccuracies in our differential gene expression analyses, we re-annotated the *melpomene* genome using the Cufflinks reference annotation-based transcript (RABT) assembly tool^33^ We used the transcriptomic data from both *H. melpomene* and *H. cydno* to reannotate the *melpomene* genome, separately for every developmental stage, and reconducted the differential gene expression analyses in DESeq2 as described above. Repeating all comparative transcriptomic analyses using these new annotations (where exons were correctly considered as part of single genes), we confirmed that both *regucalcin2* and *Grik2* were differentially expressed in both species and hybrids comparisons.

### Inference of BC3 hybrids genome composition

In order to perform comparative transcriptomic analyses between third-generation backcross hybrids (BC3) segregating at the QTL on chromosome 18 (crossing design in Supplementary figure 3), we first determined which genomic regions in these hybrids were heterozygous (*cyd/melp*) or homozygous (*cyd*/*cyd*). For this, we inferred variants from RNA-seq reads for each BC3 hybrid (individually as above), and from the combined *H. melpomene* and *H. cydno* samples. For the species, we used HaplotypeCaller^77^ on RNA-seq samples from all developmental stages of either species, to produce individual genomic records (gVCF), and then jointly genotyped *H. melpomene* and *H. cydno* gVCFs (separately for the two species) using genotypeGVCFs with default parameters. Genotype calls were filtered for quality by depth (QD) > 2, strand bias (FS) < 30 and allele depth (DP) > 4. For further analyses we kept *biallelic* genotypes only. We then used the *intersect* function of *bcftools*^72^ to infer variants exclusive to *the H. cydno* and to the *melpomene* samples.

We calculated the fraction of variants that each BC3 hybrid individual shared with the *H. melpomene* and with the *H. cydno* samples, in non-overlapping 100kb windows. We compared these to the fraction of variants that a F1 hybrid and a *H. cydno* individual (not included in the combined genotyping of the *cydno* samples), shared with the same species samples, and found that they matched either one of them, indicating heterozygous (*cyd/melp*) or homozygous (*cyd*/*cyd*) regions (Supplementary figure 5B). In this analysis, we considered only those 100kb windows where BC3 hybrids/F1 hybrid/*H. cydno* individuals shared more than 30 variants with the *melpomene*/*cydno* samples.

To corroborate our findings, we repeated the same type of analysis, this time inferring species-specific variants for *H. melpomene* and *H. cydno* using 10 *H. melpomene rosina* and 10 *H. cydno chioneus* genome resequencing samples. Variant calling files (vcf) were retrieved from Martin et al^38^. We considered only *biallelic* genotype calls that had 10 < DP < 100 and genotype quality (GQ) > 30. With this analysis we found the same heterozygous and homozygous regions in BC3 hybrids.

The size and number of the introgressed regions were in line with expectations about 3^rd^ generation backcross hybrids following our crossing design: segregating at the level of chromosome 18 and at four other chromosomes. For the BC3 hybrids sampled at 156 hours after pupal formation (APF) we had 6 *cyd/melp* and 10 *cyd*/*cyd* at the QTL region on chromosome 18 (Supplementary figure 5A), for those at 60h APF, 8 *cyd/melp* and 9 *cyd*/*cyd* hybrids at the same region. The average percentage of the genome that is heterozygous (*cyd*/*melp)* as opposed to homozygous (*cyd*/*cyd)* outside of chromosome 18 was ∼6% (close to the expectation that a 3^rd^ generation backcross genome should be 1/16 heterozygous (*cyd*/*melp*)).

### Allele-specific expression (ASE) in hybrids

In order to conduct ASE analyses we first identified species specific variants, fixed in either *H. melpomene* and *H. cydno*. For this, we took the quality filtered variants inferred from the species genome resequencing data, and assigned those genotype calls in *H. cydno* and *H. melpomene* for which allele frequency (AF) was > 0.9 as homozygous (we did not consider indels in this analysis). We then used *bcftools intersect*^73^ to get only those variants for which *H. cydno* and *H. melpomene* had opposite alleles.

At the same time, we called variants from RNA-seq reads of F1 hybrid individuals (using F1 hybrids from both datasets), again according to the GATK Best Practices (with the exception of parameters -window 35 -cluster 3, to increase SNPs density), and selected only heterozygous SNPs in F1s that matched the species-specific variants. Finally, we used GATK’s ASEReadCounter^77^, with option “-drf DuplicateRead” (without deduplicating RNA reads), to count RNA reads in the F1 hybrids (and later on in BC3 hybrids) that mapped to either the *H. cydno* or the *H. melpomene* allele. We summed all reads mapping to either the *H. cydno* or *H. melpomene* allele/variant within the same gene (both for gene models of the Hmel2.5 gene annotation and for the Cufflinks annotation we assembled previously). To test for allele specific expression (diffASE) we fitted the model “∼0 + individual + allele” in DESeq2^74^, setting library size factors to 1 (thus not normalizing between samples, as the test for diffASE is conducted within individuals). We only considered those alleles showing at least a 2-fold change in expression and *p* < 0.05, as differentially expressed.

In order to check that there were no biases in alleles assignment to one of the two species, we analyzed the ratios of the species alleles, for every gene, and checked that they were not systematically biased to either one of the two species. The log_2_ fold-changes of the species alleles were centered around 0, suggesting no obvious bias in alleles assignment^79^ (Supplementary figure 7).

### Protein-coding substitutions and predicted effects on protein-function

We inferred fixed variants in protein-coding regions from the combined *H. melpomene* and *H. cydno* RNA samples in order to include variants from genes for which we detected expression in the brain/eyes across the 3 stages. We took the quality filtered variants called from the joint genotyping of RNA-seq data of *H. cydno* and *H. melpomene* (from all stages, these did not include adult samples from the more recent sequencing batch), and selected those genotype calls for which allele frequency (AF) > 0.8, and where the allelic variant was present in at least 7 individuals of the ∼30 samples (for each species). We retained those substitutions/indels validated with the genome resequencing data. For this, of the genotype calls found in RNA reads from brain/eyes of different stages, we kept only those that were also called in at least 8 of the 10 genome resequencing samples of each species. We considered this overlapping set of variants as being fixed in *H. melpomene rosina* or *H. cydno chioneus*. Following a similar approach to Bendesky *et al*.^67^, we then restricted this set of substitutions between *H. cydno* and *H. melpomene* to protein-coding regions, and selected those non-synonymous substitutions that were considered to have moderate or high effect on protein function, with SNPeff^80^. Finally, we used the PROVEAN algorithm^35^, to further study the functional effects of these substitutions on protein function. The PROVEAN algorithm predicts the functional effect of protein sequence variations based on how they affect alignments to homologous protein sequences (for this we used the PROVEAN protein database online). We selected those amino acid changes with the suggested PROVEAN score < -2.5, indicating non-neutral effect on protein function.

### Admixture analyses

We retrieved estimated admixture proportions between *H. melpomene rosina* and *H. cydno chioneus*, for 100kb and 20kb windows, from Martin et al.^38^

## Data availability

RNA-seq data will be deposited on a public database (https://www.ebi.ac.uk/ena) on acceptance (accession ID pending).

## Code availability

Analysis scripts and behavioural data are available at: https://github.com/SpeciationBehaviour/neural_genes_heliconius.git

## Author contributions

R.M.M. and M.R. conceived the study and designed the experiments, with input from W.O.M. and C.D.J.; M.R. analysed expression and sequence data; A.E.H. analysed the behavioural data; M.R., T.J.T., S.H.M. and R.M.M. reared butterflies, dissected neural tissue and extracted RNA; R.M.M., C.D.J. and W.O.M. secured funding, contributed resources and provided supervision; R.P. additionally secured funding and contributed resources; M.R. and R.M.M. wrote the manuscript with contributions from all authors.

## Acknowledgments

We thank Chi-Yun Kuo, Liz Evans, Adriana Tapia, Moises Abanto and Morgan Oberweiser for help and assistance in the insectaries. We are grateful to Ana Pinharanda for sharing the *H. cydno* genome assembly and annotation and to Simon Martin for sharing admixture proportions and comments on the manuscript. We thank the Smithsonian Tropical Research Institute for providing research infrastructure, the Ministerio del Ambiente for permission to collect butterflies in Panama, Edinburgh Genomics and Novogene for sequencing support and Jochen Wolf who kindly provided computational resources. MR, AEH and RMM are supported by an Emmy Noether fellowship and research grant awarded to RMM by the Deutsche Forschungsgemeischaft (DFG) (Grant Number: GZ: ME 4845/1-1). RMM was also supported by a Junior Research Fellowship from King’s College Cambridge, an Ernst Mayr fellowship from the Smithsonian Tropical Research Institute, a Varley-Gradwell fellowship from the Oxford University Museum of Natural History, and the Balfour-Browne Fund, University of Cambridge. SHM was supported by a NERC IRF (NE/N014936/1). This project was also supported by the Puerto Rico Science, Technology and Research Trust (ARG 2020-00138), the Institutional Development Award (IdeA) INBRE NIH/NIGMS (award number P20 GM103475) and a National Science Foundation EPSCoR RII Track-2 FEC (award number OIA-1736026) to RP, and a European Research Council (ERC) awarded to CDJ (Grant number: 339873).

## Supplementary information

### Supplementary methods and results

#### Mapping RNA-seq reads to the *Heliconius cydno* genome

To determine whether the *H. melpomene* reference genome introduced mapping biases of RNA-seq reads, possibly affecting differential expression estimates, we also mapped to a *H. cydno* assembly/annotation^1^. Generally, we found similar patterns of differential expression when mapping to the two genomes. Since **i)** we observed an equal decrease (∼ 40 %) of genes showing 2-fold changes in *melpomene* and *cydno* when mapping to *H. cydno*, at every stage (P > 0.05 at every stage, Fisher’s Exact test, Table S2A), and **ii)** this decrease was widespread throughout the genome, we concluded that the *H. melpomene* reference genome did not bias differential gene expression analyses. We report the number of reads mapping to both genomes for each adult sample (Table S2B).

#### Allele-specific expression in the introgression line

BC3 hybrids had different combinations of chromosomes segregating for the *melpomene* alleles in a *H. cydno* background. Therefore, in principle, we could not infer *cis*- or *trans*-gene regulatory effects genome-wide from the profiles of allele specific expression (ASE) in these hybrids as for F1 hybrids, due to the diverse trans-acting environments. However, analyses (comparing gene expression levels between hybrids carrying *cyd/melp* vs. *cyd/cyd* regions on chromosomes other than 18) imply that differential expression of the candidate genes seems to be driven by the *H. melpomene* copy difference within the introgressed region on chromosome 18. Therefore, ASE analyses of candidate genes in BC3 hybrids carrying *cyd/melp* alleles on chromosome 18 should indicate whether the differences are due to cis- or trans-regulatory effects from within the introgressed region (Figure S6).

In BC3 hybrids sampled at 156h APF and 60hAPF, the *H. melpomene* and *H. cydno* alleles of the *ionotropic glutamate receptor* (*Grik2*) are expressed at very similar levels (P > 0.05 at both stages, Wald test), suggesting trans-only regulatory effects at these stages for *Grik2*. We detected diffASE expression of *regucalcin2* at 60h APF (p <0.05, Wald test), (and at 156h APF there was a tendency towards up-regulation of the *H. melpomene* allele), but at these stages *regucalcin2* was not detected as differentially expressed between pure species. Thus, although there is evidence for *cis*-regulatory effects for species differences in *regucalcin2* expression during development, the possible effect of *regucalcin2* on behaviour at these stages is less clear.

The region introgressed into *H. cydno* extended ∼3.6 Mb beyond the QTL candidate region and seven genes located within this region were differentially expressed (at either stage) in both species and backcross hybrid comparisons (Table S3). We conducted ASE analyses on these seven genes and found evidence for *cis*-regulation for only one (HMEL010030g1, p<0.001, Wald test), which had no annotated function. Unfortunately, we were unable to detect allele-informative reads for the other six genes including a serine protease inhibitor (HMEL014931g1), a CUB domain containing protein (HMEL002560g1), a methyltransferase (HMEL010030g2), a gene with a reverse transcriptase domain (HMEL034294g1), an ionotropic glutamate receptor (HMEL034304g1), and a major facilitator superfamily transporter (HMEL015745g1). (A further two genes on chromosome 18 were located outside of the introgressed region, for which we could not conduct ASE analyses because they were in *cyd*/*cyd* regions).

Our expectation was that *cis*-regulatory elements will normally act on genes in close proximity (the average distance between regulatory elements such as enhancers/repressors and the genes they regulate has been estimated to be less than 100kb^2^), making differentially expressed genes within the QTL peak – and to a lesser extent the candidate region – the best candidates. The QTL might conceivably harbour *cis*-regulatory element(s) acting on these gene(s) at a longer distance on chromosome 18 (*i*.*e*. outside of the QTL region). Although the closest is at >2.6Mb beyond the QTL peak, we currently cannot completely rule out those genes, differentially expressed in both species and backcross hybrid comparisons on chromosome 18 (Table S3), as – albeit far less well supported – candidates.

## Supplementary figures

**Figure S1.**
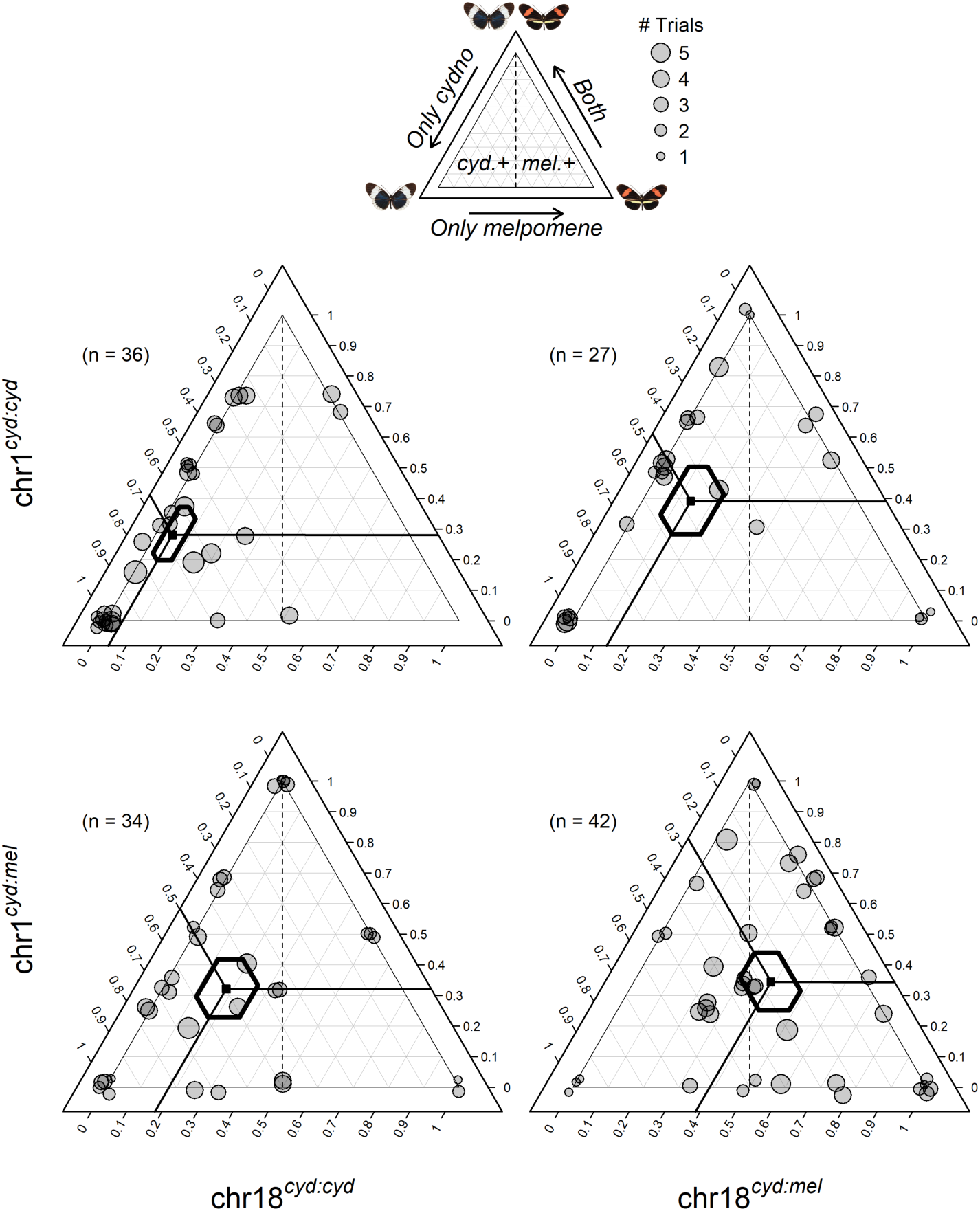
Ternary plots showing the number of 15-minute choice trials in which courtship was initiated towards *melpomene, cydno* or both females for backcross-to-*cydno* males, with different genotypes at the two QTLs retained in our model (on chromosome 1 and chromosome 18). Left ternary axis shows proportion of trials where courtship was initiated towards *H. cydno* female only, bottom axis towards *H. melpomene* female only, and right axis towards both female species. Lines project the three predicted proportions to corresponding values on the three axes and 95% credibility intervals (CrIs) for these proportions are shown as hexagons. Point size is scaled to the number of trials in which the male showed a response and a ‘jitter’ function has been applied (leading to some dots being jittered to outside the triangle).

**Figure S2.**
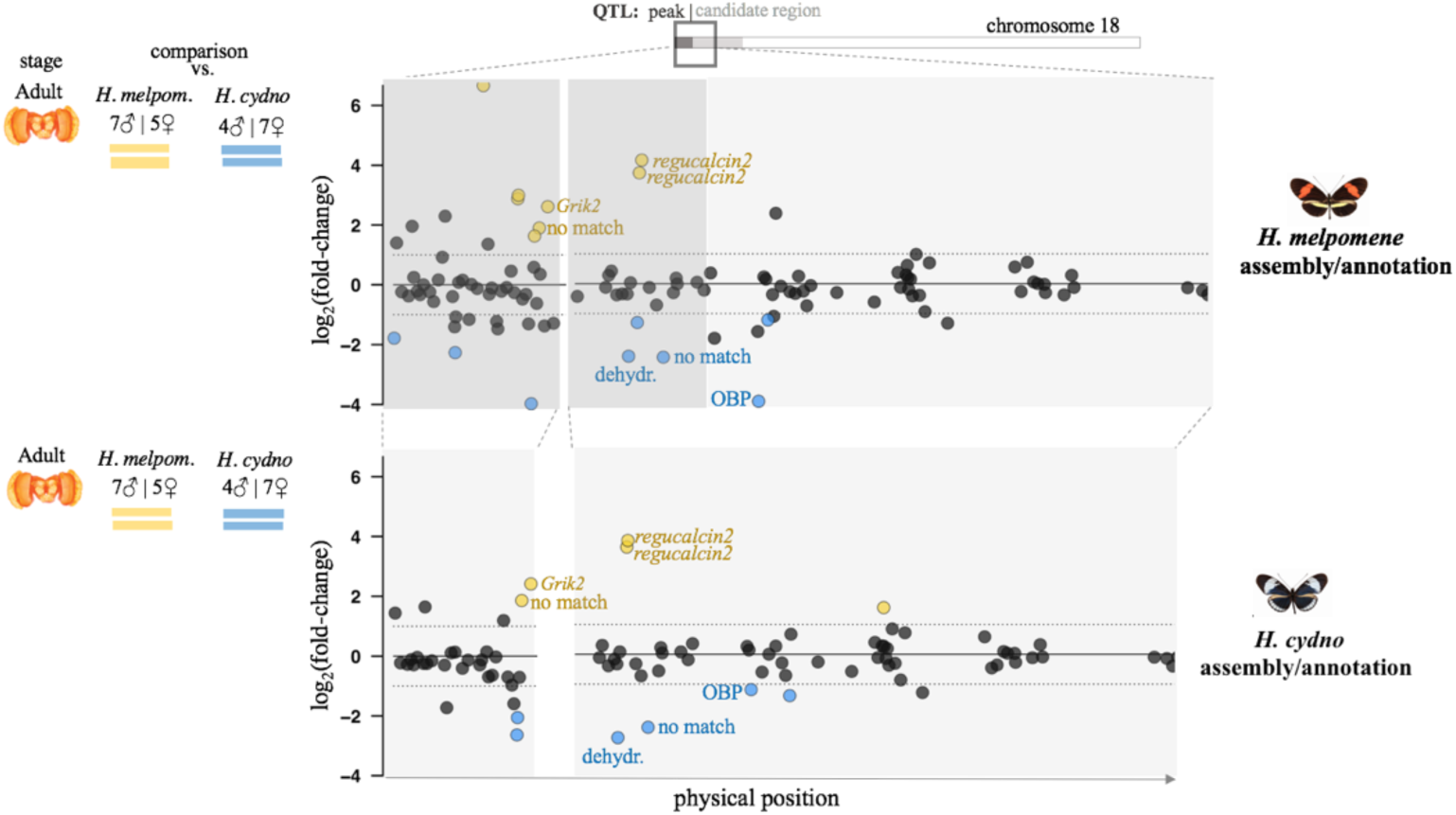
Results of comparative transcriptomic analyses between *H. melpomene* and *H. cydno* (in the imago) when mapping RNA-seq reads to the *H. melpomene* assembly/annotation (top), and to the *H. cydno* assembly/annotation (bottom), zooming in on the QTL region on chromosome 18. The *x*-axis represents physical position. Points correspond to individual genes, with the *y*-axis indicating the *log*_*2*_(fold-change) for each comparison. The two horizontal dashed lines (at *y*-values of 1 and -1) indicate a 2-fold change in expression. Genes showing a significant 2-fold+ change in expression level between groups are highlighted in orange and blue, where orange indicates higher levels in *melpomene*, blue if in *cydno*. Genes detected as differentially expressed mapping to both *melpomene* and *cydno* genomes are labelled with gene names. dehydr.=2-oxoisovalerate dehydrogenase, OBP=odorant-binding protein.

**Figure S3.**
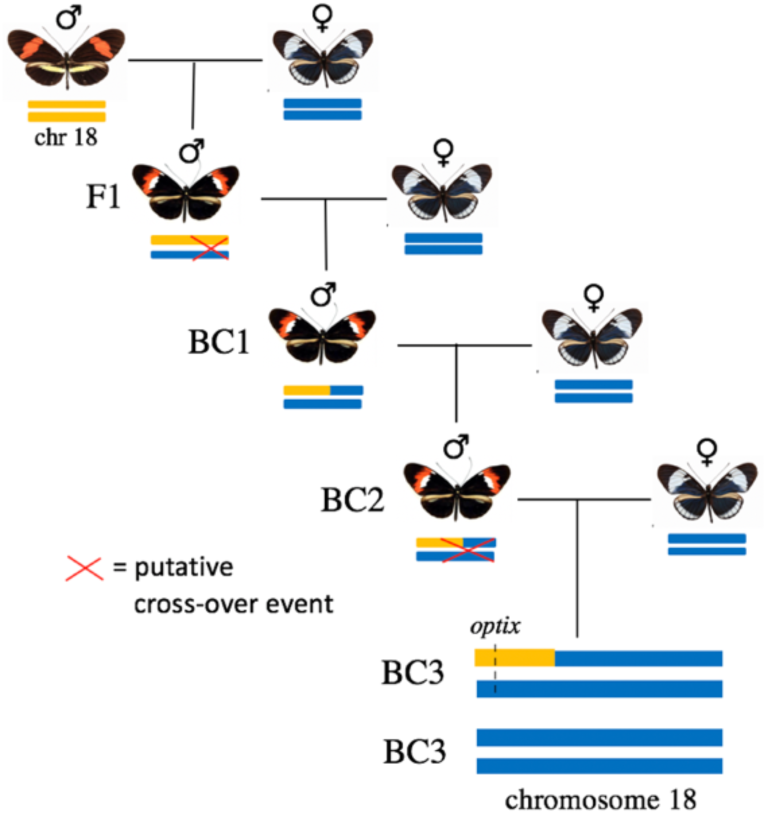
Crossing design for producing backcross hybrids segregating at the QTL on chromosome 18. This introgression line was created by outcrossing a male hybrid to *H. cydno* females over three generations, selecting a hybrid male that showed a red band on the wing at each generation. This meant that these males carried one copy of the *H. melpomene* allele at the *optix* locus. We expected that, following recombination (which occurs in males), by the fourth generation we would remain with two types of individuals: either *cyd*/*melp* or *cyd*/*cyd* at the level of the *optix* region (which approximately corresponds to the region associated with male preference behaviour).

**Figure S4.**
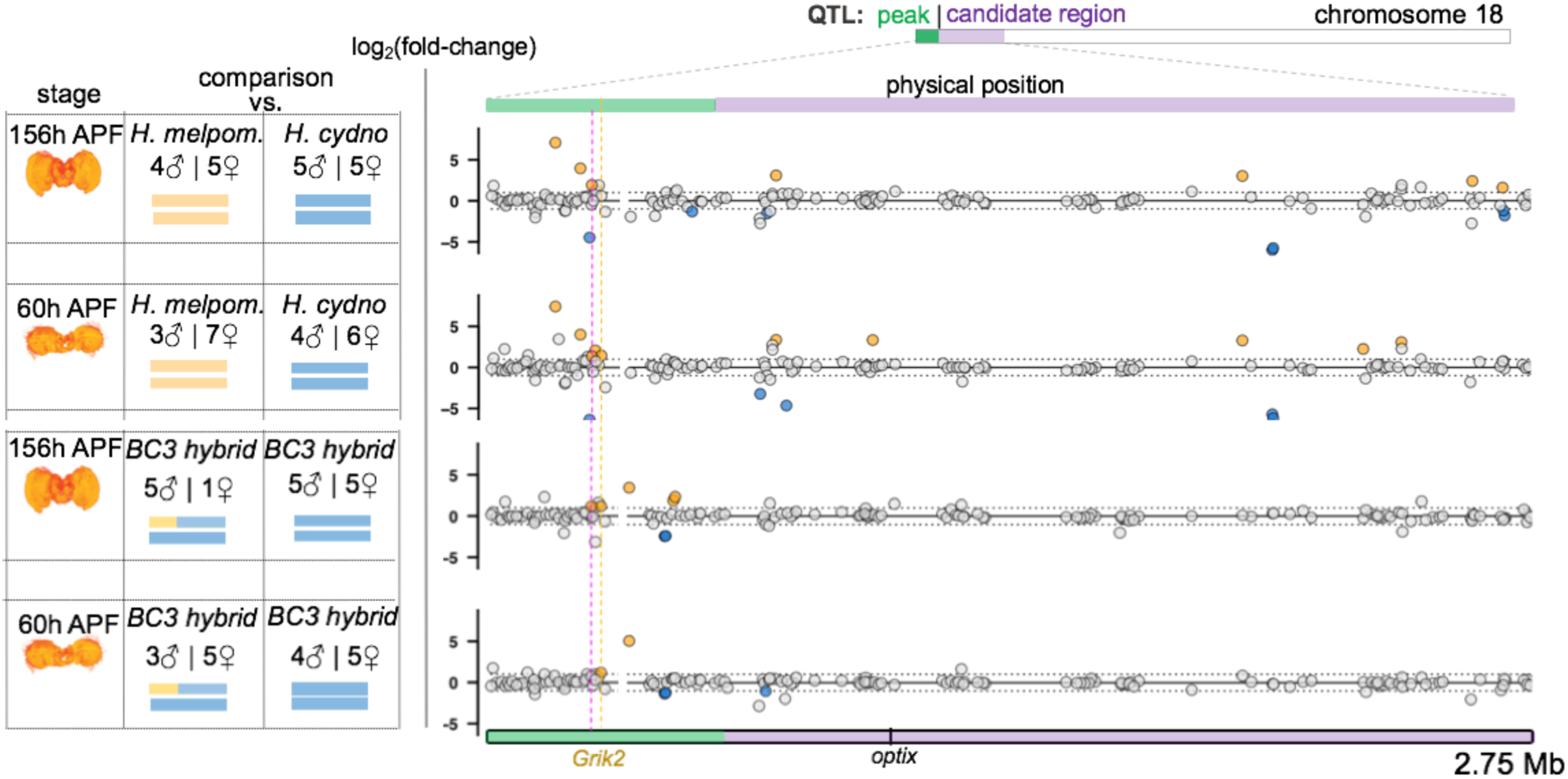
Differential gene expression at the QTL region at pupal stages. Left: summary of the comparative transcriptomic analyses with stage, number of samples and chromosome 18 composition. Right: the corresponding results, zooming in on the QTL region on chromosome 18. The *x*-axis represents physical position. The QTL peak, and the rest of the QTL 1.5 LOD candidate region are shown in green and purple, respectively. Points correspond to individual genes, with the *y*-axis indicating the *log*_*2*_(fold-change) for each comparison. The two horizontal dashed lines (at *y*-values of 1 and -1) indicate a 2-fold change in expression. Genes showing a significant 2-fold+ change in expression level between groups are highlighted in orange and blue, where orange indicates higher levels in *melpomene* or in the hybrids *cyd*/*melp* (blue if in *cydno* – hybrids *cyd*/*cyd*). Vertical dashed lines highlight those genes that are differentially expressed between *H. melpomene* and *H. cydno* AND between *cyd/melp* vs *cyd/cyd* individuals, at the same stage. One gene highlighted by a dashed fuchsia vertical line was excluded because it showed reversal of the fold change when mapping RNA-seq reads to the *H. cydno* genome.

**Figure S5.**
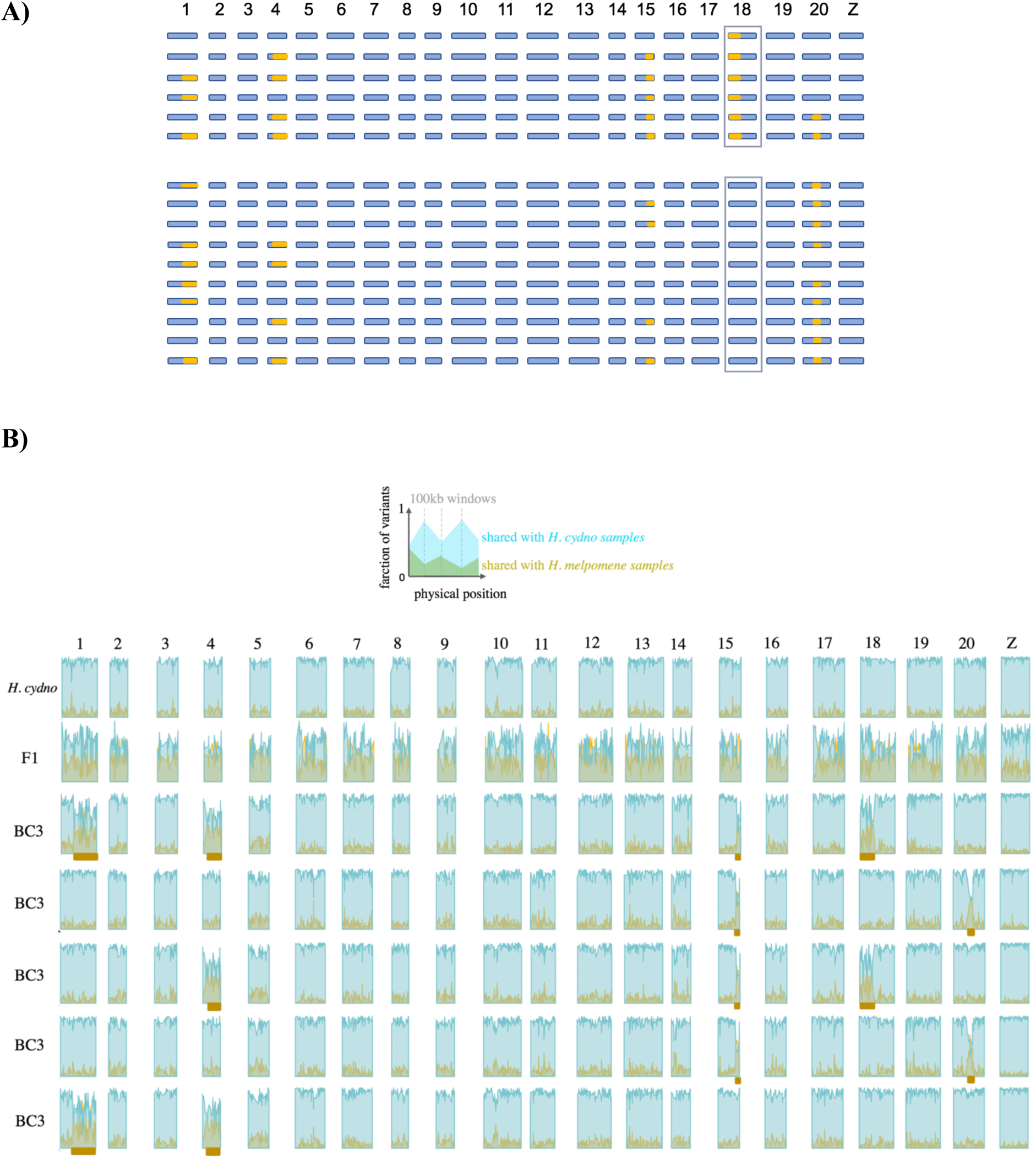
A) Schematic representation of hybrid pupae (sampled at 156h APF) genome composition. Columns represent chromosomes, rows represent individuals, orange indicates *cyd/melp* regions, blue indicates *cyd/cyd* regions. B) Genome composition of (a subset of) BC3 hybrids. We calculated the fraction of SNPs and indels that each BC3 hybrid, one *cydno* and one F1 hybrid samples shared with *H. melpomene* and *H. cydno* samples, in non-overlapping 100kb windows. x-axes represent physical position (for each chromosome), y-axes fractions of shared variants with *melpomene* (in gold) and with *cydno* (in light blue). Matching variant fractions between BC3 hybrids and the F1 hybrid, indicating heterozygous regions, are highlighted with a gold bar underneath. Note that the general trend of higher number of variants shared with *H. cydno* in heterozygous regions is due to the fact that we inferred variants by mapping to the *H. melpomene* genome (and used variant sites only for this analysis).

**Figure S6.**
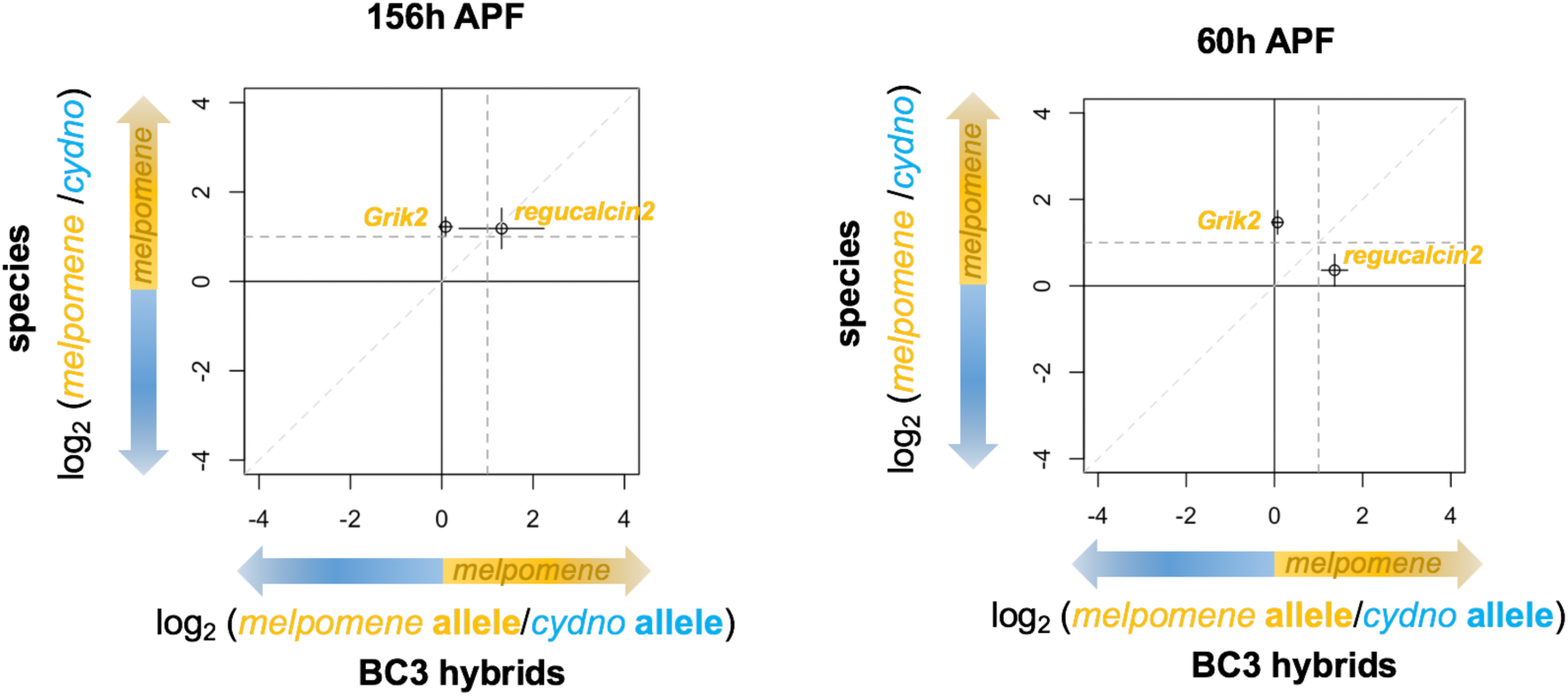
Allele specific expression profiles of candidate genes at pupal stages. Points indicate the value, and bars the standard error, of the (base 2) logarithmic fold change in expression between parental species (vertical) and the alleles in F1 hybrids (horizontal), for candidate genes (as defined in the transcript-guided annotation). Dashed lines indicate the threshold for a 2-fold change in expression for the genes in the species (horizontal), and for the alleles in the hybrids (vertical).

**Figure S7.**
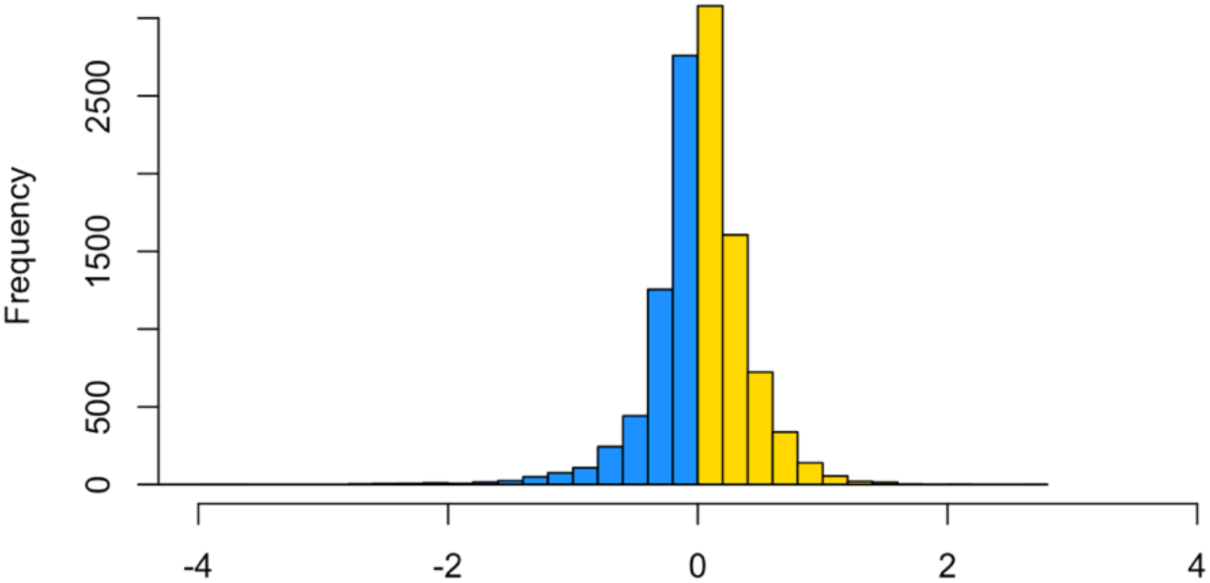
Distribution of the (base 2) logarithmic fold change in allele expression. Coloured bars indicate the number of genes showing a bias in expression for the *H. cydno* allele (in blue) and for the *H. melpomene* allele (in yellow). Values departing from 0 on the x-axis, indicate an increase in the fold change for the *H. cydno* allele (negative values) or for the *H. melpomene* allele (positive values), respectively.

## Supplementary tables

**Table S1.**
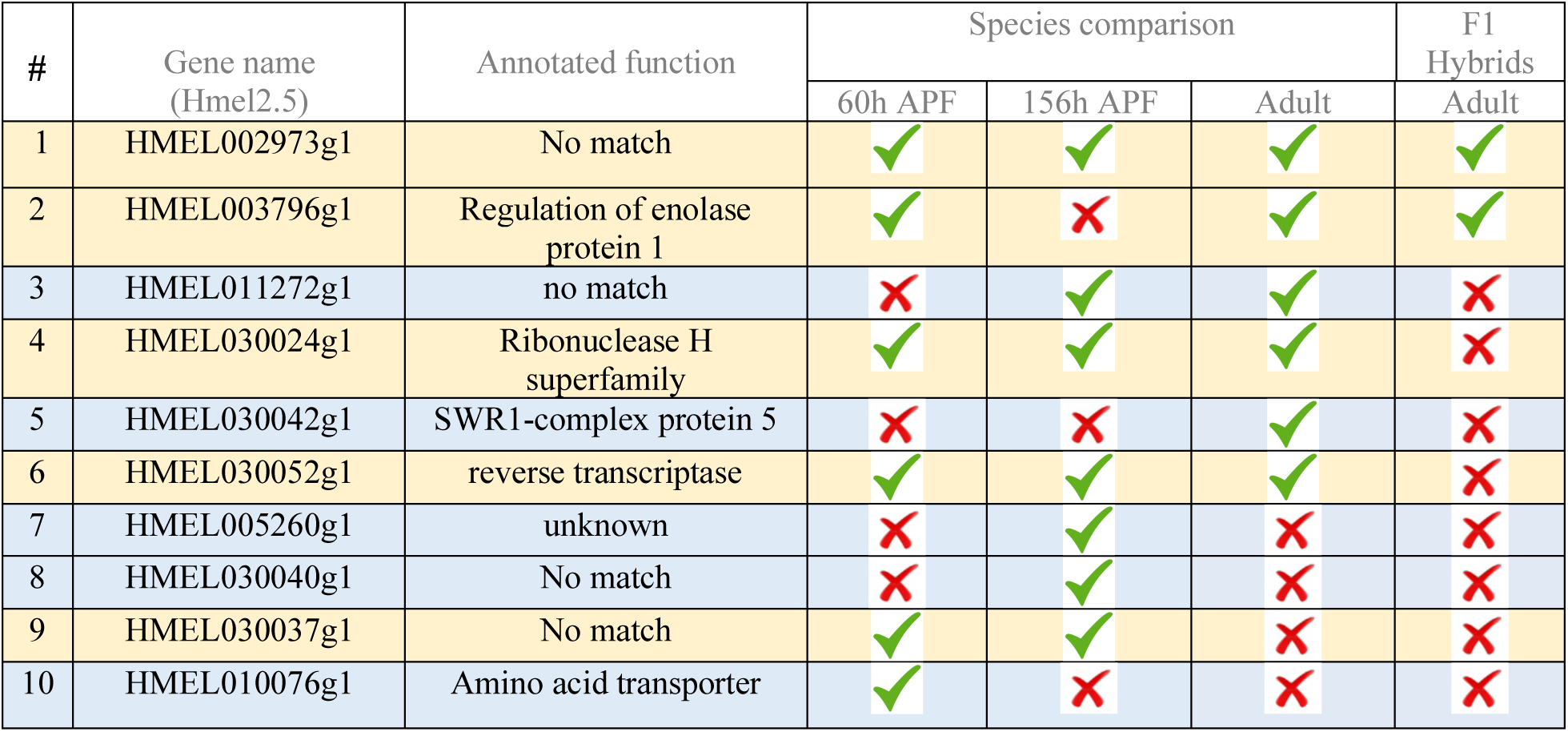
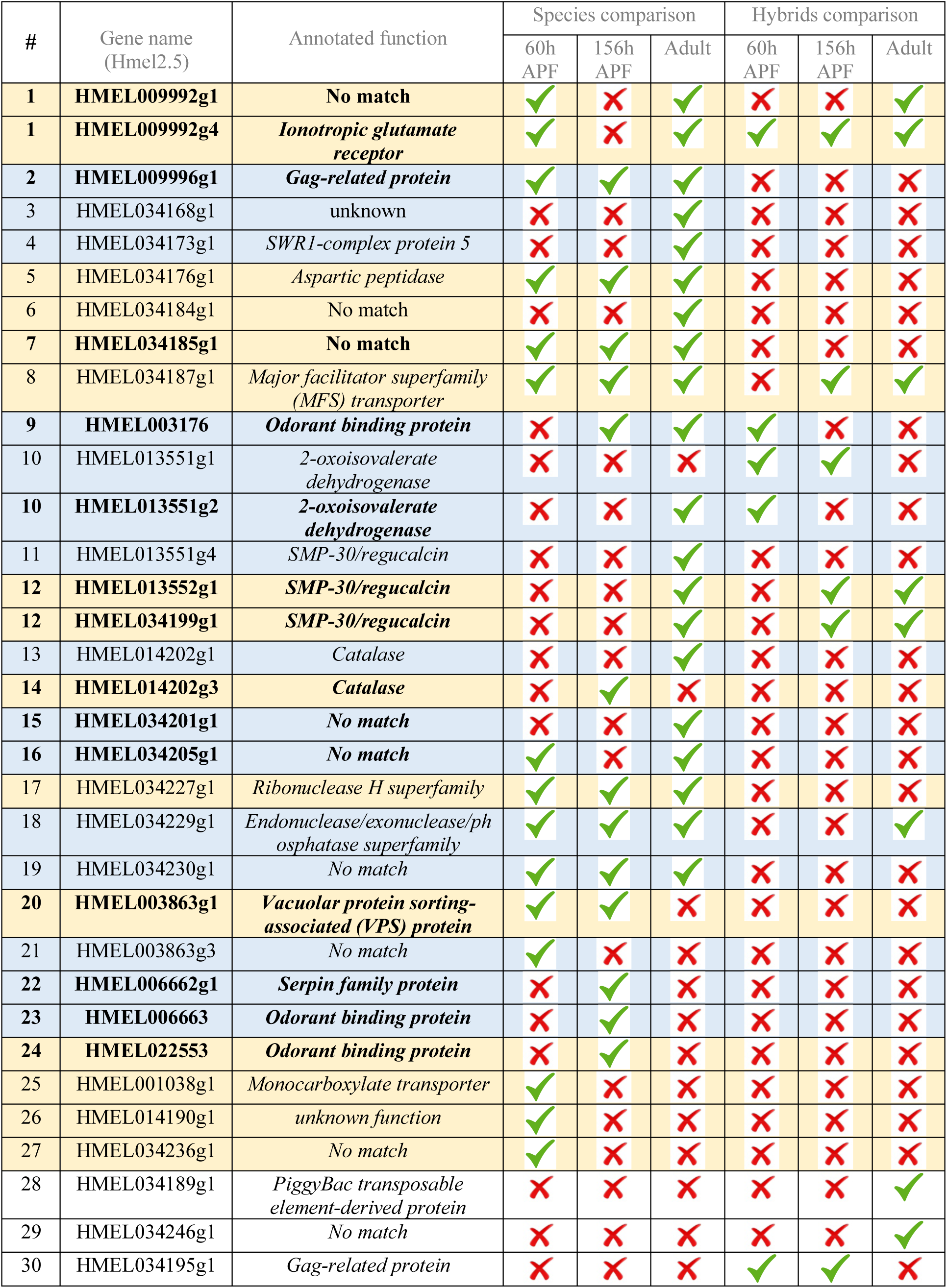
List of differentially expressed genes in species and hybrids comparisons. **A)** QTL chromosome 1. Orange indicates genes up-regulated in *H. melpomene*, and blue those up-regulated in *H. cydno*. **B)** QTL chromosome 18. Those genes found to be differentially expressed when also mapping to the *H. cydno* genome are highlighted in bold. Genes annotated as distinct but sharing the same number in the table (#) were later found to be single genes (see second paragraph of the Results section).

| # | Gene name<br>(Hmel2.5) | Annotated function | Species comparison |  |  | F1<br>Hybrids |
| --- | --- | --- | --- | --- | --- | --- |
|  |  |  | 60h APF | 156h APF | Adult | Adult |
| 1 | HMEL002973g1 | No match | ✓ | ✓ | ✓ | ✓ |
| 2 | HMEL003796g1 | Regulation of enolase<br>protein 1 | ✓ | ✗ | ✓ | ✓ |
| 3 | HMEL011272g1 | no match | ✗ | ✓ | ✓ | ✗ |
| 4 | HMEL030024g1 | Ribonuclease H<br>superfamily | ✓ | ✓ | ✓ | ✗ |
| 5 | HMEL030042g1 | SWR1-complex protein 5 | ✗ | ✗ | ✓ | ✗ |
| 6 | HMEL030052g1 | reverse transcriptase | ✓ | ✓ | ✓ | ✗ |
| 7 | HMEL005260g1 | unknown | ✗ | ✓ | ✗ | ✗ |
| 8 | HMEL030040g1 | No match | ✗ | ✓ | ✗ | ✗ |
| 9 | HMEL030037g1 | No match | ✓ | ✓ | ✗ | ✗ |
| 10 | HMEL010076g1 | Amino acid transporter | ✓ | ✗ | ✗ | ✗ |

| # | Gene name<br>(Hmel2.5) | Annotated function | Species comparison |  |  | Hybrids comparison |  |  |
| --- | --- | --- | --- | --- | --- | --- | --- | --- |
|  |  |  | 60h<br>APF | 156h<br>APF | Adult | 60h<br>APF | 156h<br>APF | Adult |
| 1 | **HMEL009992g1** | **No match** | ✓ | ✗ | ✓ | ✗ | ✗ | ✓ |
| 1 | **HMEL009992g4** | ***Ionotropic glutamate receptor*** | ✓ | ✗ | ✓ | ✓ | ✓ | ✓ |
| 2 | **HMEL009996g1** | ***Gag-related protein*** | ✓ | ✓ | ✓ | ✗ | ✗ | ✗ |
| 3 | HMEL034168g1 | unknown | ✗ | ✗ | ✓ | ✗ | ✗ | ✗ |
| 4 | HMEL034173g1 | *SWR1-complex protein 5* | ✗ | ✗ | ✓ | ✗ | ✗ | ✗ |
| 5 | HMEL034176g1 | *Aspartic peptidase* | ✓ | ✓ | ✓ | ✗ | ✗ | ✗ |
| 6 | HMEL034184g1 | No match | ✗ | ✗ | ✓ | ✗ | ✗ | ✗ |
| 7 | **HMEL034185g1** | **No match** | ✓ | ✓ | ✓ | ✗ | ✗ | ✗ |
| 8 | HMEL034187g1 | *Major facilitator superfamily (MFS) transporter* | ✓ | ✓ | ✓ | ✗ | ✓ | ✓ |
| 9 | **HMEL003176** | ***Odorant binding protein*** | ✗ | ✓ | ✓ | ✓ | ✗ | ✗ |
| 10 | HMEL013551g1 | *2-oxoisovalerate dehydrogenase* | ✗ | ✗ | ✗ | ✓ | ✓ | ✗ |
| 10 | **HMEL013551g2** | ***2-oxoisovalerate dehydrogenase*** | ✗ | ✗ | ✓ | ✓ | ✗ | ✗ |
| 11 | HMEL013551g4 | *SMP-30/regucalcin* | ✗ | ✗ | ✓ | ✗ | ✗ | ✗ |
| 12 | **HMEL013552g1** | ***SMP-30/regucalcin*** | ✗ | ✗ | ✓ | ✗ | ✓ | ✓ |
| 12 | **HMEL034199g1** | ***SMP-30/regucalcin*** | ✗ | ✗ | ✓ | ✗ | ✓ | ✓ |
| 13 | HMEL014202g1 | *Catalase* | ✗ | ✗ | ✓ | ✗ | ✗ | ✗ |
| 14 | **HMEL014202g3** | ***Catalase*** | ✗ | ✓ | ✗ | ✗ | ✗ | ✗ |
| 15 | **HMEL034201g1** | **No match** | ✗ | ✗ | ✓ | ✗ | ✗ | ✗ |
| 16 | **HMEL034205g1** | **No match** | ✓ | ✗ | ✓ | ✗ | ✗ | ✗ |
| 17 | HMEL034227g1 | *Ribonuclease H superfamily* | ✓ | ✓ | ✓ | ✗ | ✗ | ✗ |
| 18 | HMEL034229g1 | *Endonuclease/exonuclease/phosphatase superfamily* | ✓ | ✓ | ✓ | ✗ | ✗ | ✓ |
| 19 | HMEL034230g1 | No match | ✓ | ✓ | ✓ | ✗ | ✗ | ✗ |
| 20 | **HMEL003863g1** | ***Vacuolar protein sorting-associated (VPS) protein*** | ✓ | ✓ | ✗ | ✗ | ✗ | ✗ |
| 21 | HMEL003863g3 | No match | ✓ | ✗ | ✗ | ✗ | ✗ | ✗ |
| 22 | **HMEL006662g1** | ***Serpin family protein*** | ✗ | ✓ | ✗ | ✗ | ✗ | ✗ |
| 23 | **HMEL006663** | ***Odorant binding protein*** | ✗ | ✓ | ✗ | ✗ | ✗ | ✗ |
| 24 | **HMEL022553** | ***Odorant binding protein*** | ✗ | ✓ | ✗ | ✗ | ✗ | ✗ |
| 25 | HMEL001038g1 | *Monocarboxylate transporter* | ✓ | ✗ | ✗ | ✗ | ✗ | ✗ |
| 26 | HMEL014190g1 | unknown function | ✓ | ✗ | ✗ | ✗ | ✗ | ✗ |
| 27 | HMEL034236g1 | No match | ✓ | ✗ | ✗ | ✗ | ✗ | ✗ |
| 28 | HMEL034189g1 | *PiggyBac transposable element-derived protein* | ✗ | ✗ | ✗ | ✗ | ✗ | ✓ |
| 29 | HMEL034246g1 | No match | ✗ | ✗ | ✗ | ✗ | ✗ | ✓ |
| 30 | HMEL034195g1 | *Gag-related protein* | ✗ | ✗ | ✗ | ✓ | ✓ | ✗ |

**Table S2.** **A)** Number of genes showing significant >2-fold change in expression, at different stages, mapping to the *H. melpomene* and to the *H. cydno* genomes. Note that the considerable reduction in the number of genes detected as differentially expressed when mapping to *H. cydno* is most likely a result of the lower quality/completeness of the *H. cydno* genome assembly. **B)** Number of reads mapping to the *H. melpomene* and the *H. cydno* genome for every species sample at the adult stage.

| Stage | Mapping to: | Up-regulated in *H. melpomene* | Up-regulated in *H. cydno* |
| --- | --- | --- | --- |
| Adult | *H. melpomene* | 694 | 733 |
|  | *H. cydno* | 390 | 451 |
| 156h APF | *H. melpomene* | 837 | 667 |
|  | *H. cydno* | 518 | 403 |
| 60h APF | *H. melpomene* | 846 | 642 |
|  | *H. cydno* | 490 | 376 |

| *H. melpomene* |  |  | *H. cydno* |  |  |
| --- | --- | --- | --- | --- | --- |
| ID | mapped to *H. mel* | mapped to *H. cyd* | ID | mapped to *H. mel* | mapped to *H. cyd* |
| 45 | 14660303 | 14399385 | 50 | 17617312 | 10276127 |
| 47 | 7157775 | 7106443 | 51 | 14965350 | 11157953 |
| 53 | 14013806 | 13804267 | 57 | 17460091 | 10958479 |
| 70 | 11850181 | 11656033 | 58 | 20474961 | 14651415 |
| 71 | 11218116 | 11023267 | 67 | 27415308 | 15742957 |
| 78 | 14114255 | 13618292 | 68 | 22159665 | 12175286 |
| 80 | 14551439 | 14097670 | 81 | 34947798 | 19085351 |
| 83 | 13616160 | 13260160 | 82 | 15525206 | 10264627 |
| 100 | 13771837 | 13458527 | 84 | 13139892 | 9836625 |
| 104 | 12802894 | 12664628 | 98 | 18846622 | 12585701 |
| 128 | 13840154 | 13489851 | 99 | 20309368 | 13271219 |
| 218 | 16942305 | 16812556 |  |  |  |

**Table S3.** List of differentially expressed genes in both species and backcross hybrid comparisons (at 60h APF or 156h APF). Genes annotated as distinct but sharing the same number in the table (#) were later found to be single genes. *gene excluded because it showed reversal of the fold change when mapping to the *H. cydno* genome.

| # | Gene name (Hmel2.5) | Annotated function | 60h APF | 156h APF | chromosome | Start (bp) position | End (bp) position |
| --- | --- | --- | --- | --- | --- | --- | --- |
| 1 | HMEL009992g4 | *Ionotropic glutamate receptor* | ✓ | ✗ | 18 | 334856 | 345144 |
| * | HMEL034187g1 | *no match* | ✗ | ✓ | 18 | 314734 | 315069 |
| 2 | HMEL014931g1 | Serine endopeptidase inhibitor | ✓ | ✓ | 18 | 3260635 | 3264491 |
| 3 | HMEL034294g1 | Reverse transcriptase domain | ✓ | ✓ | 18 | 4369133 | 4370180 |
| 4 | HMEL002560g1 | CUB domain | ✓ | ✗ | 18 | 4679193 | 4684506 |
| 5 | HMEL034304g1 | *Ionotropic glutamate receptor* | ✓ | ✗ | 18 | 4935938 | 4944768 |
| 6 | HMEL010030g2 | *Methyltransferase* | ✓ | ✗ | 18 | 5503482 | 5505157 |
| 6 | HMEL010030g3 | *no match* | ✓ | ✗ | 18 | 5506399 | 5508526 |
| 7 | HMEL010030g1 | *no match* | ✓ | ✓ | 18 | 5496842 | 5498456 |
| 8 | HMEL015745g1 | *Major facilitator superfamily* | ✗ | ✓ | 18 | 5976551 | 5982484 |
| 9 | HMEL014795g1 | *Major facilitator superfamily* | ✗ | ✓ | 18 | 10194935 | 10200873 |
| 10 | HMEL015842g1 | *Alpha crystallin/heat shock protein* | ✗ | ✓ | 18 | 13546671 | 13547216 |
| 11 | HMEL030024g1 | *Ribonuclease H superfamily* | ✗ | ✓ | 1 | 1009751 | 1013056 |

**Table S4.** Heterozygosity on the *Z*-chromosome. Heterozygosity is calculated as proportion of variants (SNPs and indels) which are heterozygous, in each sample, rounded at the second decimal place (note that variant sites were inferred having mapped to the *H. melpomene* genome).

| *H. melpomene* |  |  |  | *H. cydno* |  |  |  |
| --- | --- | --- | --- | --- | --- | --- | --- |
| Males |  | Females |  | Males |  | Females |  |
| ID | Het. | ID | Het. | ID | Het. | ID | Het. |
| Adults |  |  |  |  |  |  |  |
| 45 | 0.49 | 53 | 0.04 | 57 | 0.23 | 50 | 0.02 |
| 47 | 0.46 | 78 | 0.05 | 82 | 0.23 | 51 | 0.02 |
| 70 | 0.47 | 80 | 0.04 | 98 | 0.25 | 58 | 0.02 |
| 71 | 0.48 | 128 | 0.04 | 99 | 0.24 | 67 | 0.02 |
| 83 | 0.46 | 218 | 0.05 |  |  | 68 | 0.01 |
| 100 | 0.49 |  |  |  |  | 81 | 0.02 |
| 104 | 0.46 |  |  |  |  | 84 | 0.02 |
| 156h APF |  |  |  |  |  |  |  |
| 5 | 0.47 | 6 | 0.05 | 4 | 0.26 | 13 | 0.02 |
| 14 | 0.49 | 18 | 0.05 | 8 | 0.26 | 21 | 0.04 |
| 17 | 0.49 | 24 | 0.05 | 142 | 0.30 | 30 | 0.03 |
| 184 | 0.50 | 150 | 0.05 | 151 | 0.29 | 137 | 0.04 |
|  |  | 220 | 0.06 | 156 | 0.29 | 168 | 0.04 |
| 60h APF |  |  |  |  |  |  |  |
| 92 | 0.48 | 87 | 0.06 | 85 | 0.26 | 90 | 0.02 |
| 97 | 0.49 | 95 | 0.05 | 86 | 0.27 | 118 | 0.02 |
| 115 | 0.47 | 117 | 0.05 | 119 | 0.27 | 125 | 0.02 |
|  |  | 124 | 0.04 | 146 | 0.26 | 144 | 0.02 |
|  |  | 149 | 0.06 |  |  | 162 | 0.02 |
|  |  | 164 | 0.05 |  |  | 200 | 0.02 |
|  |  | 208 | 0.06 |  |  |  |  |

| Hybrids |  |  |  |  |  |  |  |
| --- | --- | --- | --- | --- | --- | --- | --- |
| Males |  | Females |  | Males |  | Females |  |
| ID | Het. | ID | Het. | ID | Het. | ID | Het. |
| F1 hybrids (adults) |  |  |  |  |  |  |  |
| 42 | 0.64 | 56 | 0.05 |  |  |  |  |
| 49 | 0.64 | 69 | 0.04 |  |  |  |  |
| Introgression line -156h APF |  |  |  | Introgression line – 60h APF |  |  |  |
| 105 | 0.26 | 108 | 0.02 | 152 | 0.27 | 161 | 0.02 |
| 116 | 0.27 | 123 | 0.02 | 193 | 0.27 | 165 | 0.02 |
| 126 | 0.27 | 136 | 0.02 | 198 | 0.25 | 179 | 0.02 |
| 131 | 0.27 | 139 | 0.02 | 199 | 0.27 | 187 | 0.02 |
| 133 | 0.26 | 140 | 0.02 | 201 | 0.26 | 188 | 0.02 |
| 154 | 0.26 | 166 | 0.02 | 212 | 0.28 | 192 | 0.02 |
| 155 | 0.26 |  |  | 215 | 0.27 | 197 | 0.02 |
| 183 | 0.27 |  |  |  |  | 209 | 0.03 |
| 185 | 0.27 |  |  |  |  | 214 | 0.02 |
| 189 | 0.27 |  |  |  |  | 224 | 0.02 |

